# Simulations of the effect of diffusion on asymmetric spin echo based quantitative BOLD: An investigation of the origin of deoxygenated blood volume overestimation

**DOI:** 10.1101/570697

**Authors:** Alan J Stone, Naomi C Holland, Avery J L Berman, Nicholas P Blockley

## Abstract

Quantitative BOLD (qBOLD) is a technique for mapping oxygen extraction fraction (OEF) and deoxygenated blood volume (DBV) in the human brain. Recent measurements using an asymmetric spin echo (ASE) based qBOLD approach produced estimates of DBV which were systematically higher than measurements from other techniques. In this study, we investigate two hypotheses for the origin of this DBV overestimation using simulations and consider the implications for experimental measurements. Investigations were performed by combining Monte Carlo simulations of extravascular signal with an analytical model of the intravascular signal.

**Hypothesis 1:** DBV overestimation is due to the presence of intravascular signal which is not accounted for in the analysis model. Intravascular signal was found to have a weak effect on qBOLD parameter estimates.

**Hypothesis 2:** DBV overestimation is due to the effects of diffusion which are not accounted for in the analysis model. The effect of diffusion on the extravascular signal was found to result in a vessel radius dependent variation in qBOLD parameter estimates. In particular, DBV overestimation peaks for vessels with radii from 20 to 30 μm and is OEF dependent. This results in the systematic underestimation of OEF.

**Implications:** The impact on experimental qBOLD measurements was investigated by simulating a more physiologically realistic distribution of vessel sizes with a small number of discrete radii. Overestimation of DBV consistent with previous experiments was observed, which was also found to be OEF dependent. This results in the progressive underestimation of the measured OEF. Furthermore, the relationship between the measured OEF and the true OEF was found to be dependent on echo time and spin echo displacement time.

The results of this study demonstrate the limitations of current ASE based qBOLD measurements and provide a foundation for the optimisation of future acquisition approaches.

## Introduction

The quantitative BOLD (qBOLD) technique is a relaxometry based approach for mapping oxygen extraction fraction (OEF) and deoxygenated blood volume (DBV) in the human brain (He and Yablonskiy, 2007). An elevated OEF is indicative of tissue at risk of infarction, such as the penumbral tissue surrounding the core infarct of an ischaemic stroke (Astrup et al., 1981). When combined with a measurement of cerebral blood flow (CBF), the cerebral metabolic rate of oxygen consumption (CMRO_2_) can also be estimated (Kety and Schmidt, 1948). Since qBOLD can provide this valuable information in a non-invasive and rapidly acquired manner, it has a great deal of potential for providing these quantitative physiological measurements in clinical research applications.

The analytical model used to analyse qBOLD data assumes that the signal decay behaves as though it were in the static dephasing regime (SDR) i.e. the diffusion of water in tissue does not influence the signal decay due to magnetic field inhomogeneity (Yablonskiy and Haacke, 1994). However, simulations of the Gradient Echo Sampling of Spin Echo (GESSE) pulse sequence, which is often used to acquire qBOLD data, have shown that this is not the case and that diffusion introduces a vessel size dependent effect on the signal decay (Dickson et al., 2010; Pannetier et al., 2014). However, qBOLD data can also be acquired using the Asymmetric Spin Echo (ASE) pulse sequence, which provides a direct measurement of the reversible relaxation rate, R_2_′, and eliminates the need to remove R_2_-weighting from the acquired signal (required by GESSE) (An and Lin, 2003; Stone and Blockley, 2017). Nevertheless, it is unclear whether a similar diffusion effect is present in ASE data. Interestingly estimates of DBV made using this ASE based acquisition are systematically higher than those reported for GESSE based measurements (He and Yablonskiy, 2007), suggesting that different effects may be at play.

The overestimation of DBV by ASE based qBOLD is at least partially responsible for the underestimation of the OEF (Stone and Blockley, 2017). This overestimation has previously been suggested to be due to the presence of intravascular blood signal, which is not accounted for in the analytical qBOLD model, with flow crushing gradients proposed as a solution (An and Lin, 2003). However, since it has been shown that diffusion results in additional signal attenuation (Dickson et al., 2010), which is similarly unaccounted for in the analytical qBOLD model, this may also provide a mechanism for DBV overestimation.

In this study, we investigate both mechanisms to discover whether either can account for the overestimation of DBV in ASE based qBOLD. The effect of diffusion on the extravascular tissue signal was examined using Monte Carlo simulations (Boxerman et al., 1995) and the intravascular blood signal was simulated using a recently published analytical model (Berman and Pike, 2018). Whilst these effects are initially considered using simulations with vessels of a single radius, these results are also integrated using a more physiologically realistic vessel size distribution to investigate sources of systematic error in real world measurements.

## Theory

Transverse signal decay results from dephasing of the net magnetisation due to the presence of magnetic field inhomogeneity at multiple scales. The effect of these scales on the qBOLD signal can be considered with reference to a spin echo pulse sequence. At the microscopic scale spins experience local magnetic field inhomogeneities caused by neighbouring spins that are rapidly varying. Due to this rapid magnetic field variation, the phase evolution cannot be rewound by the application of a refocussing pulse. The resulting signal decay is described by the irreversible transverse relaxation rate R_2_. The macroscopic scale describes magnetic field inhomogeneity on the scale of the head e.g. due to the nasal sinuses or ear canals. This effect can be reversed by a refocussing pulse due to its static nature, enabling the phase evolution in the period before the refocussing pulse to be rewound by the time the spin echo is formed. At this scale the signal decay is described by the reversible relaxation rate R_2_′ and referred to as the SDR. At intermediate scales, often referred to as the mesoscopic scale, diffusion becomes increasingly important as the so called Diffusion Narrowing Regime is approached. The transition between these regimes is dependent on the scale of the magnetic field inhomogeneity and the distance the spin travels due to diffusion. More precisely, the characteristic diffusion time, τ_D_,

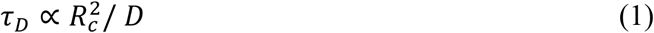

which is dependent on the radius, R_c_, of the deoxygenated blood vessel and the diffusion coefficient, D, is on the order of the time taken by a water molecule to diffuse a distance equivalent to the radius of the vessel (Yablonskiy and Haacke, 1994). This results in an averaging of the magnetic field distribution surrounding the vessels and a loss of phase history, meaning that signal cannot be efficiently recovered by a refocussing pulse.

### Modelling the qBOLD signal

The qBOLD model relies on the known relationship between R_2_′ and the baseline OEF, E_0_, and deoxygenated blood volume fraction, *V*_0_, for a network of randomly oriented blood vessels approximated as infinite cylinders (Yablonskiy and Haacke, 1994),

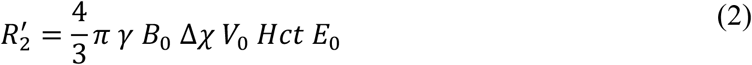

where γ is the proton gyromagnetic ratio, *B*_0_ is the main magnetic field, Δ_*χ*_ is the difference in volume magnetic susceptibility between fully oxygenated and fully deoxygenated blood in CGS units and *Hct* is the haematocrit. In this work it was assumed that the arterial oxygen saturation is 100% and hence E_0_=1-Y, where Y is the venous oxygen saturation. Further modelling has shown that the R_2_′-weighted signal is not purely monoexponential and can be approximated for two distinct regimes (An and Lin, 2000; Yablonskiy and Haacke, 1994),

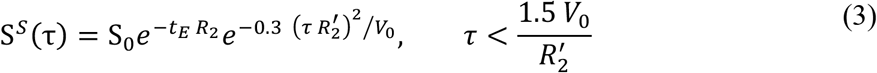

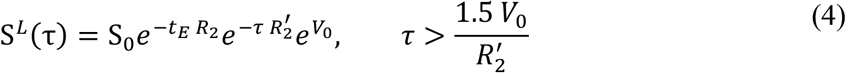

where *t*_E_ is the echo time and τ is the spin echo displacement time. In the long τ regime (S^*L*^ Eq. (4)) the signal decay takes a monoexponential form, whilst in the short τ regime (S^*S*^, Eq. (3)) the signal follows a quadratic exponential form. A log-linear fit to long τ data enables *R*_2_′ to be estimated. Furthermore, comparison of the measured signal at 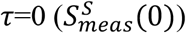 with the intercept extrapolated from long τ data 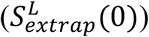 enables *V*_0_ to be calculated.

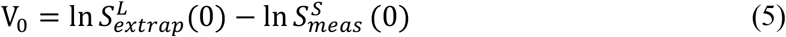

Henceforth we will refer to this as the SDR qBOLD model.

### Simulating the effect of diffusion

Monte Carlo simulations of the qBOLD signal were performed by repeating the following three steps for each simulated proton.

Step 1: Generate a system of vessels. The vessel system was constrained to fit within a sphere of radius *R*_*s*_. Vessel origin points (*O*) were randomly selected, with half placed on the surface of the sphere and half within the sphere to ensure a homogenous vessel density following previous work (Dickson et al., 2010). A uniform distribution of points over the surface of the sphere was ensured by generating a unit vector (*X*_*i*_) from a normally distributed random number generator (mean 0, standard deviation 1) and scaling by *R*_s_ (Muller, 1959). Within the sphere, uniform density was maintained by taking account of the increased volume occupied by points far from the centre of the system. This scaling factor, *U*, is selected from a uniform distribution of random numbers (range 0 to 1).

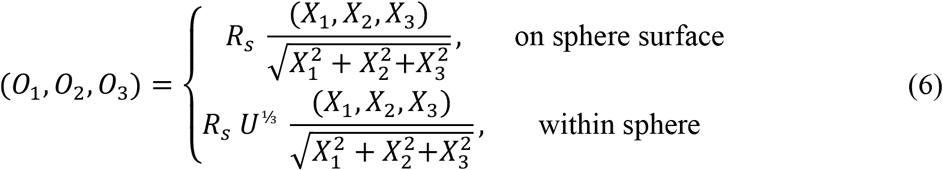

Vessels were modelled as randomly oriented infinitely long cylinders with a single radius, R_c_, placed at the vessel origin points described by Eq. (6) and extended out to the surface of the sphere. This enable the volume occupied by each vessel to be calculated with further vessels added to the system until the target blood volume fraction (*V*_f_) was reached. Random orientation was ensured by generating a unit vector from a normally distributed random number generator (mean 0, standard deviation 1).

Step 2: Proton random walk. Protons are initially placed at the centre of the vessel system. Each step taken by the proton is independently selected along each dimension from a normal distribution of random numbers with mean 0 and standard deviation m with diffusion coefficient, D, and time interval between steps, Δ*t*.

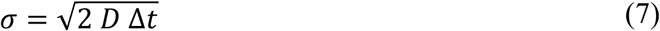

Step 3: Estimate the phase accrued at each step. The phase, Δϕ, accumulated by the proton during each time interval is calculated by summing over the field contributions from all *N* vessels (Boxerman et al., 1995),

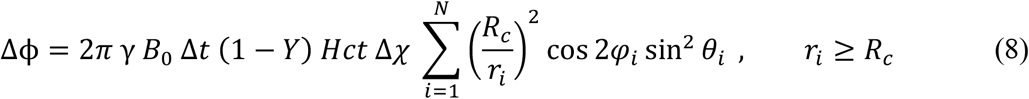

where *θ* is the angle of the vessel with respect to *B*_0_, *φ* is the angle with respect to the projection of *B*_0_ onto a plane orthogonal to the vessel, *r*_*i*_ is the perpendicular distance to the vessel and *Y* is the blood oxygen saturation (see Fig. 1). Only the equation for the magnetic field outside of the vessel is presented, since only extravascular signal was simulated.

By appropriate combination of the phase accrued in each interval it is possible to simulate the phase evolution of the ASE and GESSE pulse sequences as a function of τ, ϕ(τ).

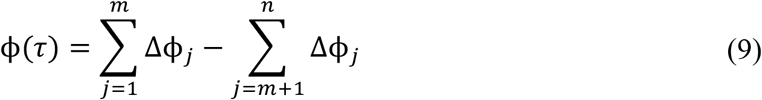

where *m* defines the transition from signal decay to signal recovery due to the refocussing pulse, *n* is the point at which the signal is acquired and where 0 ≤ *m* ≤ *n*. For ASE *m* = (*t*_*E*_ − *τ*) / 2 ∆*t* and *n* = *t*_*E*_ / ∆*t*, whilst for GESSE *m* = *t*_*SE*_ / 2 ∆*t* and *n* = (*t*_*SE*_ + τ) / ∆*t*. Here, *t*_*E*_ is defined as the timing of the centre of the readout and *t*_*SE*_ is the time at which the spin echo forms (see Fig. 2). These definitions reflect an important distinction between the ASE and GESSE pulse sequences, whereby *t*_*E*_ is fixed for ASE and variable for GESSE whilst *t_SE_* is variable for ASE and fixed for GESSE.

**Fig. 1.**
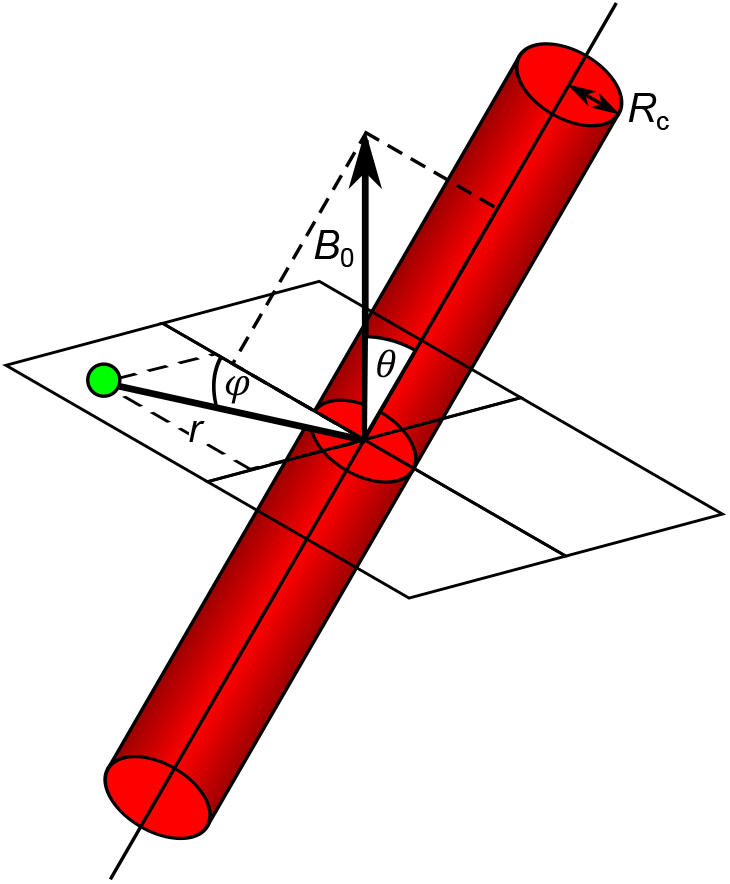
Blood vessels are approximated as infinitely long cylinders at an angle *θ* with respect to *B*_0_, the main magnetic field. The proton location is defined to be on a plane orthogonal to the blood vessel at a perpendicular distance *r* and an angle ϕ with respect to the projection of *B*_0_ onto the plane.

The phase evolution of *P* protons is then summed to simulate the decay of the extravascular ASE or GESSE signal (Weisskoff et al., 1994),

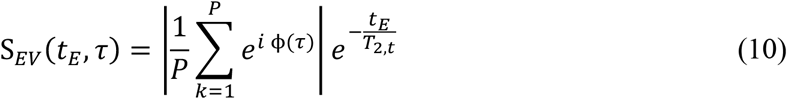

where *T*_2*,t*_ is the underlying tissue T_2_.

**Fig. 2.**
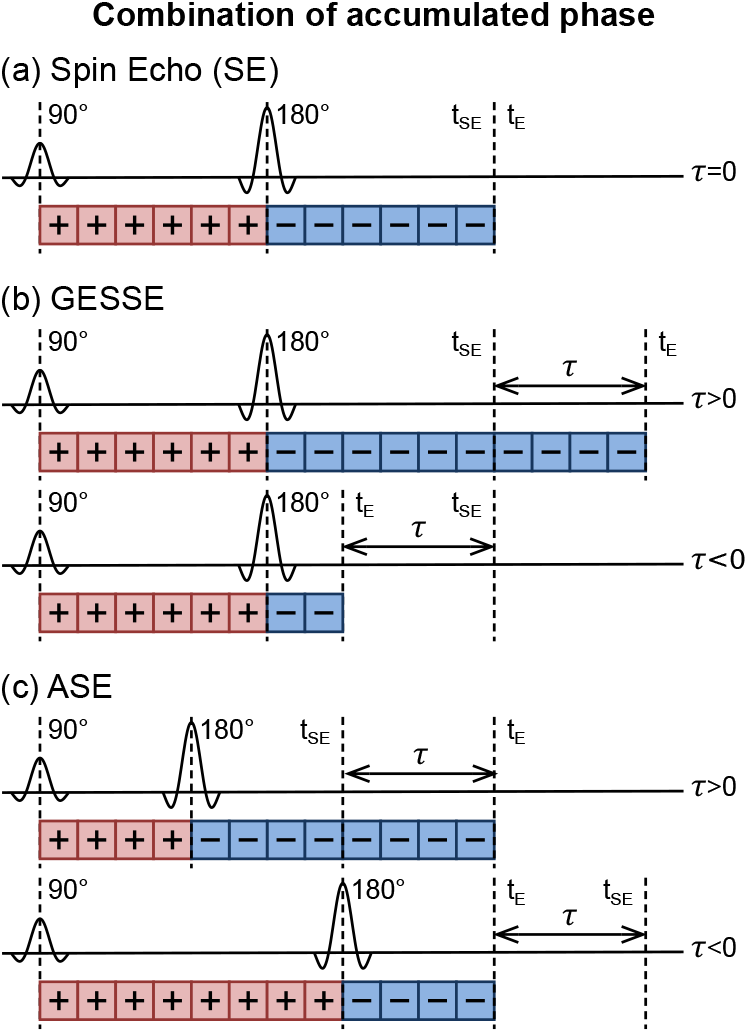
By combining the phases generated by the Monte Carlo simulations different pulse sequences can be simulated. The cumulative sum of the phase accumulated after the 180° refocussing pulse is subtracted from the cumulative sum of the phases accumulated prior to the refocussing pulse. In a standard spin echo pulse sequence (a) the refocussing pulse is placed midway between the 90° excitation pulse and the echo time (t_E_), which is equal to the spin echo time (t_SE_). The GESSE pulse sequence (b) introduces R_2_′-weighting through the parameter τ by altering t_E_, whilst keeping t_SE_ constant. Note: each value of τ is acquired at a different t_E_. The ASE sequence (c) introduces R_2_′-weighting by shifting the refocussing pulse by a time τ/2 leading to a change in t_SE_, although t_E_ is kept constant. By convention positive values of τ occur when the t_E_>t_SE_ and negative values occur when t_E_<t_SE_.

Intravascular signal has traditionally been difficult to simulate, with empirical measurements of blood R_2_ and R_2_^*^ commonly used (Griffeth and Buxton, 2011). However, simulating the R_2_′-weighted signal using the difference between R_2_ and R_2_^*^ is likely to be inaccurate in the short τ regime. Recently an analytical model of the blood signal during a Carr-Purcell Meiboom-Gill (CPMG) pulse sequence was extended to capture the signal evolution between an arbitrary number of spin echoes (Berman and Pike, 2018), i.e. the conditions that exist for ASE and GESSE pulse sequences. Using this model, the intravascular signal, *S*_*IV*_, is described by,

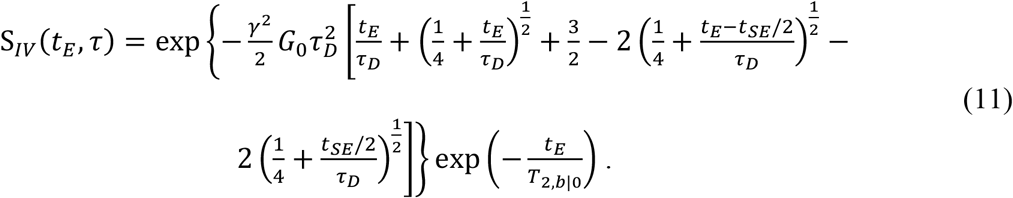

Here τ_D_=R_rbc_^2^/D_b_, where R_rbc_ is the characteristic size of red blood cells and D_b_ is the diffusion coefficient of blood, *T*_2,*b*|0_ is the intrinsic T_2_ of blood (measured when the blood is fully oxygenated) and G_0_ is the mean square field inhomogeneity in blood (Berman et al., 2017),

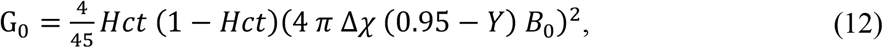

where the value of 0.95 represents the red blood cell oxygen saturation which is equal to the susceptibility of plasma (Spees et al., 2001). The value of *t*_*SE*_ is fixed for GESSE but is variable for ASE with *t*_*SE*_ = *t*_E_ − τ. By definition *t*_*E*_ is fixed for ASE and varying for GESSE.

Finally, the total signal, *S*_*TOT*_, is calculated by taking a volume weighted sum of the intra- and extravascular signals.

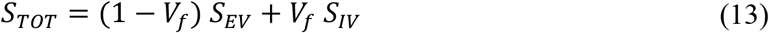

It should be noted that if the simulated blood vessels contain deoxygenated blood then *V*_f_ is equivalent to *V*_0_, the deoxygenated blood volume.

## Methods

### Simulations

Simulations of the tissue signal were performed following the theory outlined above. Firstly, extravascular signal decay was simulated using Monte Carlo simulations (B_0_=3 T, γ=267.5×10^6^ rad s^−1^ T^−1^). The radius of the spherical system of vessels, *U*, was chosen to maintain a similar number of vessels, *N*, regardless of the vessel radius (*N*~1,300). For each proton, a complete random walk was generated with a step size, Δ*t*, of 20 μs, which was downsampled to 200 μs, and *D*=1 μm^2^ms^−1^ (Boxerman et al., 1995). The perpendicular distance, *r*_*i*_, to each vessel in the system was then calculated. For protons that passed close to vessels, defined as 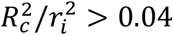, the perpendicular distance was recalculated using the original 20 μs time step to better sample the rapid magnetic field variation expected close to vessels (Dickson et al., 2010). Walks that moved the proton inside a vessel were flagged to be discarded in order to simulate non-permeable blood vessels, rather than reflecting the proton at the vessel surface which is less computationally efficient. This approach does not prevent protons passing close to vessels (as defined above) and due to the reduced time step used under this condition the spatial variation in the magnetic field is well sampled. The phase of each proton was allowed to evolve for 120 ms after the excitation with Δχ=0.27 ppm in CGS units (Spees et al., 2001). Phase accrual was stored for each proton in 2 ms intervals, Δ*t*. A new system of vessels was generated for each proton and a total of 10,000 protons were simulated for each vessel radius investigated. However, the number of protons that passed within a vessel increased as vessel radius was reduced (26% at 5 μm versus 3.1% at 1 mm at *V*_f_=3%). Therefore, only the first P=5,000 protons that did not pass within a vessel were used to calculate *S*_*EV*_ using Eq. (10) with *T*_2,*t*_ =80 ms. Secondly, intravascular signal decay was simulated using Eqs. (11) and (12), which are independent of vessel radius. Based on previous work the following parameters were used (Berman et al., 2017): *T*_2,*b*|0_ =189 ms, R_rbc_ =2.6 μm and D_b_ =2 μm^2^ms^−1^. The total signal was then calculated using Eq. (13).

Whilst the intravascular simulations are rapid to perform, Monte Carlo simulations of the extravascular signal are time consuming. Therefore, the following approaches were taken to accelerate these simulations, with examples presented as supplementary figures. We have previously shown that different oxygenation levels can be simulated by scaling the accrued phase of a nominal oxygenation value by the target value (Blockley et al., 2008). This is made possible by saving the phase of each proton and the fact that phase is a linear function of blood oxygenation for a network of vessels with the same oxygenation (Fig. S1a). Different volume fractions can be simulated from the signal magnitude generated by Eq. (10). It has been shown that the extravascular signal, *S*_*EV*_, can be described as a radius dependent shape function, *f*(*R*_*c*_, τ), scaled by the volume fraction (Dickson et al., 2011; Kiselev and Posse, 1999) (Fig. S2a).

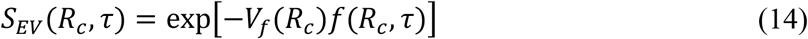

It is also possible to simulate the effect of different rates of diffusion using the results of existing Monte Carlo simulations. Since the effect on the signal decay is dependent on the characteristic diffusion time, τ_D_, then Eq. (1) provides an alternative way of simulating a change in the diffusion coefficient. For example, the signal simulated from vessels with R_c_=5 μm and D=1 μm^2^ms^−1^ is equivalent to the signal produced by simulations with R_c_=7 μm and D=2 μm_2_ms^−1^ i.e. doubling D requires R_c_ to be increased by 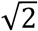 (Fig. S3a). Since the diffusion coefficient is expected to vary in the range 0.78 to 1.09 μm^2^ms^−1^ in cortical grey matter (Helenius et al., 2002), this is equivalent to between an 11.6% reduction and a 4.4% increase in vessel radius. As such, the diffusion coefficient wasn’t varied in the following simulations, relying on variation in R_c_ to examine the range of characteristic diffusion times.

Finally, it is possible to simulate the effect of a system with multiple vessel radii by combining multiple single vessel radius simulations of the extravascular signal (Dickson et al., 2011; Kiselev and Posse, 1999). The resulting combined signal, 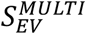, can be calculated as the product of the signals of M single vessel simulations which have already been scaled for blood oxygenation and volume fraction as described above (Fig. S4a).

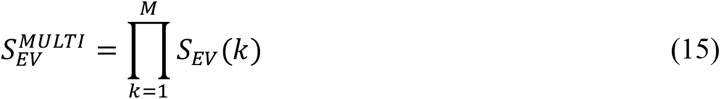

The total signal including the contribution from intravascular blood can then be calculated using Eq. (13). In this case the blood volume fraction is only equivalent to DBV when the vessel distribution does not include fully oxygenated blood vessels.

When combined these acceleration approaches vastly reduce simulation time. The average duration of a Monte Carlo simulation for a single vessel radius was 2 hrs 25 mins. In contrast, scaling existing Monte Carlo results takes on the order of 100 ms. This enables new investigations to be performed which were previously prohibitively time consuming. Analysis of the fidelity of signals generated by scaling existing simulations versus direct simulation showed that in general the percentage error ((*S*^*simulated*^ − *S*^*scaled*^)/*S*^*simulated*^) is less than 2% (Fig. S1b, S2b, S3b and S4b).

### Parameter quantification

The following framework was used to quantify the parameters of the qBOLD model from the simulated decay curves. The parameters of the SDR qBOLD model (*R*_2_′ and DBV) were organised as a vector of unknowns (x) in a linear system (A⋅x=B) (Stone et al., 2019). The first row of the matrix A represents Eq. (3) when τ=0 with subsequent rows representing Eq. (4) with values of τ beyond the transition between the quadratic and linear exponential regime. In this case only values of τ greater than 15 ms were used to be consistent with previous qBOLD experiments (Stone and Blockley, 2017). Vector B contains the ASE signals, *S*(*τ*).

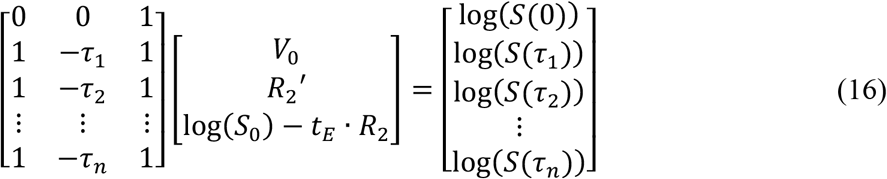

Parameters were estimated via Eq. (16) using the least square solution, with the error in each parameter determined from the covariance matrix. Finally, OEF can be estimated by rearranging Eq. (2).

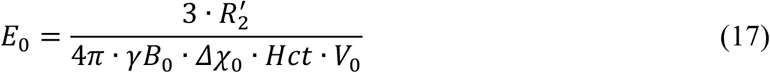

### Effect of diffusion on ASE signal decay

Initial simulations were performed for a selection of vessel radii (R_c_=5, 10, 50, 1000 μm), a venous Y of 60%, a Hct of 40% and a DBV of 3%. Simulations of the ASE pulse sequence were performed with t_E_=60 ms and −60 ms ≤ τ ≤ 60 ms for both extra- and intravascular signal, where τ =60 ms corresponds to pure gradient echo decay. For validation purposes, similar simulations were performed for the GESSE pulse sequence using t_SE_=60 ms and −30 ms ≤ τ ≤ 60 ms. Hence a common t_E_/t_SE_ was chosen to be consistent with previous simulations (Dickson et al., 2010).

### Effect of diffusion on qBOLD parameters

A further set of synthetic ASE signal decay curves were generated for vessel radii logarithmically spaced between 1 and 1,000 μm. All other parameters were set consistent with previous experimental qBOLD measurements (Stone and Blockley, 2017). In the context of these simulations this required *t*_*E*_=80 ms with *τ*=0 and *τ*=16 to 64 ms in 4 ms steps (Δτ). The *apparent* value of R_2_′, DBV and OEF were then estimated using Eq. (16) and (17). The effect of diffusion on the estimation of qBOLD parameters was investigated by first fixing OEF and varying DBV and then by fixing DBV and varying OEF. In the former case a fixed OEF of 40% was coupled with DBV values of 1, 3 and 5%, whilst in the latter case DBV was fixed at 3% and OEF took values of 20, 40 and 60%. These values are considered to be the *true* parameters in both cases. The results of varying DBV were also used to consider the percentage error in DBV as a function of vessel radius i.e. 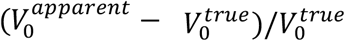. In these single vessel simulations arterial blood is assumed to have an oxygen saturation, *Y*, of 100% hence the venous oxygen saturation, *Y*_*ν*_ = 1 − *E*_0_.

The effect of intravascular signal on qBOLD parameter estimates was investigated by repeating these simulations, but excluding the intravascular compartment. In this way it was possible to quantify the percentage of the parameter estimate (PE) which results from the presence of intravascular signal i.e. (*PE*_*EV*_ − *PE*_*EV+IV*_)/*PE*__*EV+IV*__.

Further investigation of the effect of diffusion on DBV estimates was pursued based on a consideration of Eq. (5), which suggests that errors must be due to either the signal measured at 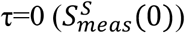 or the extrapolated estimate of the signal at τ=0 from the R_2_′ fit 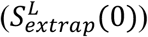, or both. However, given the analysis represented by Eq. (16), 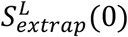 is not estimated and 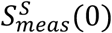 is confounded by T_2_ decay. The latter was corrected by calculating the signal decay relative to the value at R=1,000 μm, where previous simulations would suggest the SDR applies and hence signal attenuation should be zero. The former was estimated by subtracting this relative measure of 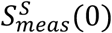 from the estimated value of apparent DBV.

### Effect of a physiologically realistic vessel radius distribution

The effect of a more physiologically realistic distribution of vessel radii was investigated by integrating the results from single radius simulations. A compartmental model of the vasculature derived from the morphology of the sheep brain was selected (Sharan et al., 1989). This model has five orders of arterial and venous vessels, with a range of radii, and a capillary compartment with a single vessel radius (Table 1). Additional Monte Carlo simulations for this range of vessel radii were performed and combined using the acceleration techniques described above. Arterial vessels were assigned an arterial oxygen saturation, Y_a_, of 98%, which was used to calculate the venous saturation, Y_v_, for a given OEF.

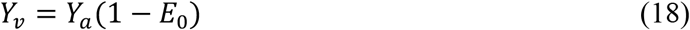

**Table 1.**
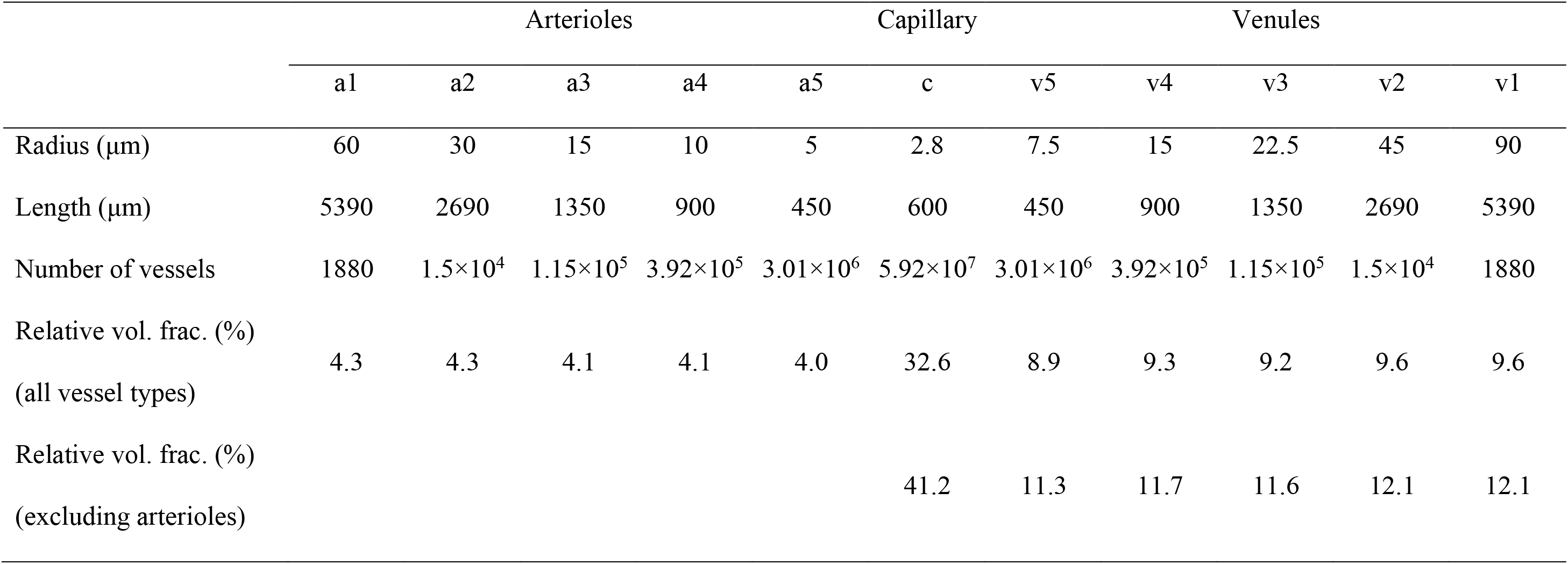
Vascular compartment model described by (Sharan et al., 1989). Radius, length and number of vessels were used to calculate the relative volume fractions for each compartment with and without arteriolar vessels.

The capillary compartment was an intermediate oxygen saturation, Y_c_, calculated as an average of the arterial and venous saturations weighted by a factor, κ, equal to 0.4 representing a weighting towards the venous saturation (Griffeth and Buxton, 2011; Tsai et al., 2003).

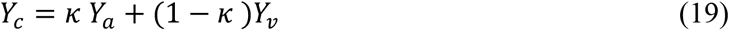

Relative blood volume fractions for each vessel type were calculated by estimating the volume of each vessel radius population as cylinders with the properties described in Table 1. These relative blood volume fractions were then scaled by the total cerebral blood volume (CBV). Pairs of OEF and CBV values were drawn from a uniform random number generator within the following ranges: OEF 0-100%, CBV 0-10%. The qBOLD parameters were quantified for 1,000 random OEF-CBV pairs to examine the effect of diffusion across the physiological range. In the absence of a strict definition of DBV, the ground truth was assumed to be equal to the combined blood volume occupied by capillary and venous vessels. This is therefore only a working assumption, since it is likely the *true* DBV is weighted by blood oxygenation and vessel radius. Deoxyhaemoglobin content, dHb, was calculated based on the same assumption for DBV and a value for the density of brain tissue ρ=1.04 g/ml (Rempp et al., 1994) using the following equation.

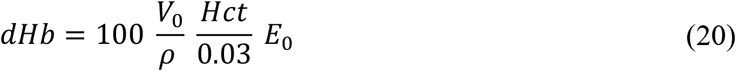

For comparison these simulations were also repeated for the original ASE based qBOLD implementation with t_E_=64 ms with *τ*=0 and *τ*=10 to 18 ms in 4 ms steps (An and Lin, 2003).

Details on how to access the simulation code, simulation results and analysis code that underlie this study can be found in Appendix A.

## Results

Figure 3 presents simulations of the signal generated by the ASE pulse sequence in the absence of T_2_ decay and with an initial transverse magnetisation of one at *t*_E_ =0. The extravascular signal (Fig. 3a) was found to be symmetric with respect to the spin echo (τ=0) regardless of vessel radius. Similarly, the intravascular signal (Fig. 3b) was symmetric, but displayed a relatively weak signal decay as a function of τ. In contrast, simulations of the GESSE pulse sequence demonstrated increasing asymmetry with reducing vessel radius for the extravascular signal and strong asymmetry for the intravascular signal (Fig. S5).

**Fig. 3.**
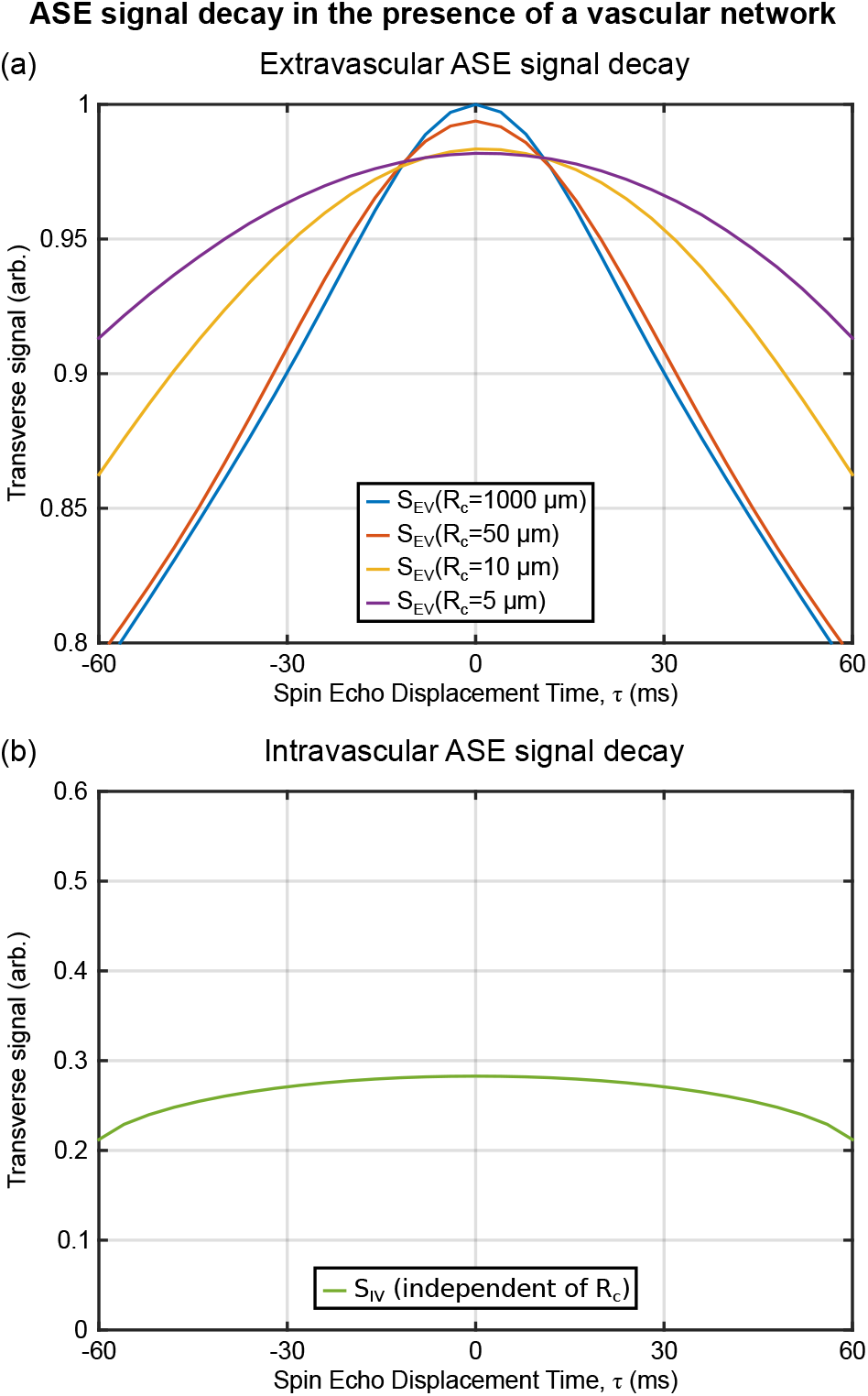
Examples of the signal decay from the ASE pulse sequence as a function of vessel radius (Y=60%, V_0_=3%, Hct=40%). (a) The extravascular signal (S_EV_) decay is observed to be symmetric with respect to τ=0 regardless of vessel radius. Signal attenuation at τ=0 increases as vessel radius decreases due to the increased effect of diffusion. (b) The intravascular signal (S_IV_) decay shows considerable signal attenuation which is symmetric and varies weakly with τ.

Figure 4 displays the effect of vessel size on the parameter estimates from the SDR qBOLD model. The apparent values of R_2_′ plateau above a critical vessel radius of approximately 40 μm (Fig. 4a,d) and are then consistent with predictions from the SDR qBOLD model (dashed lines calculated using Eq. (2)). The apparent DBV is found to be strongly dependent on vessel radius, peaking between 20 and 30 μm (Fig. 4b,e). Estimates of the apparent OEF increase monotonically with vessel radius reaching the value predicted by the SDR qBOLD model as the vessel radius approaches 1,000 μm (Fig. 4c,f). When the true OEF was fixed whilst DBV was varied (Fig. 4c) estimates of apparent OEF were consistent across DBV levels, suggesting that the error in DBV is a linear scale factor. Likewise, it can be seen that the profile of apparent DBV when the true DBV was fixed and OEF was varied (Fig. 4e) peak at different vessel radius values, suggesting that the error in DBV is OEF dependent. Furthermore, this effect can be seen to result in a reduced dynamic range for the estimates of apparent OEF as vessel size is reduced (Fig. 4f). Figure 5 confirms that the percentage error in DBV is constant for a given combination of OEF and vessel radius (Fig. 5a), but differs for different OEF values (Fig. 5b). For reference, an increase in the value of the diffusion coefficient would result in a linear translation to the right along the x-axis for data plotted against such a log vessel radius.

**Fig. 4.**
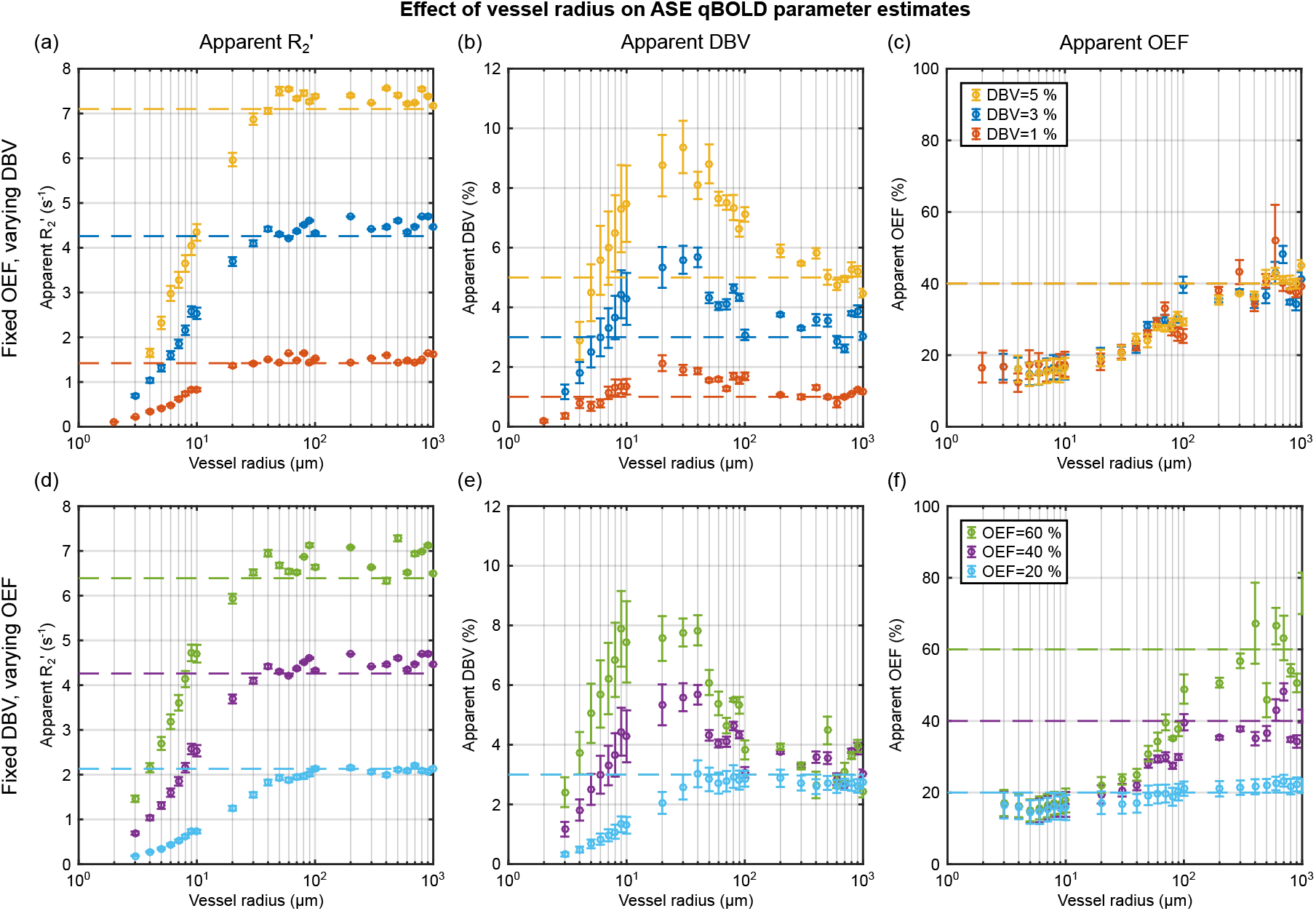
Investigation of the effect of vessel radius on the parameter estimates derived from ASE based qBOLD. Simulations were first performed with a fixed OEF (E_0_=40%) and three DBV values (top) then with a fixed DBV (V_0_=3%) and three values of OEF (bottom). The apparent R_2_′ (left) is estimated for each OEF-DBV pair and presented alongside the R_2_′ values predicted by the SDR qBOLD model (dashed lines). Likewise, the apparent DBV (centre) and apparent OEF (right) are presented alongside the true DBV and OEF (dashed lines).

**Fig. 5.**
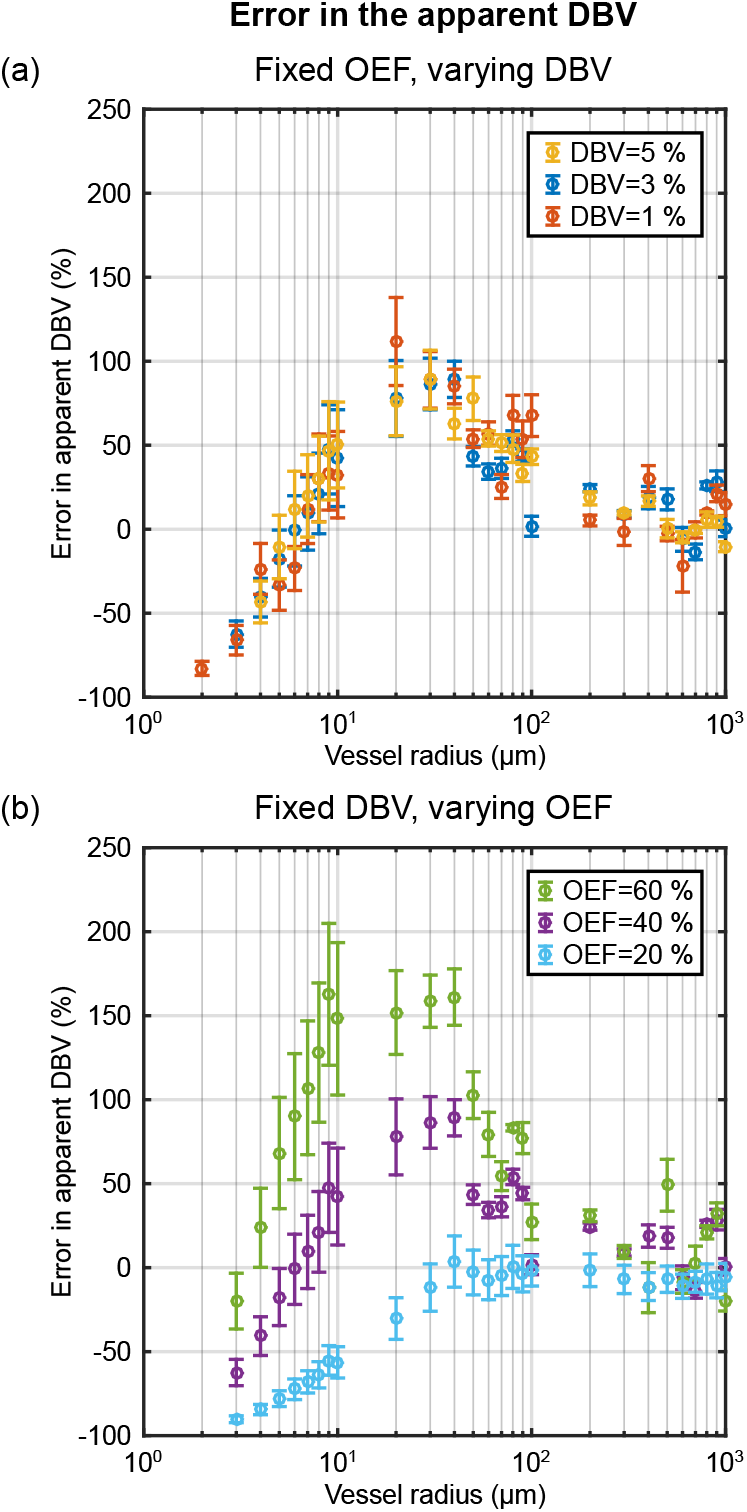
Estimation of the percentage error in DBV at each of the three simulated values with (a) fixed E_0_=40%, varying DBV and (b) fixed V_0_=3%, varying E_0_. The largest error is observed for vessels between 10 and 100 μm and is smallest as vessel size approaches 1 mm. The magnitude of the error is also dependent on OEF. The percentage error was calculated as 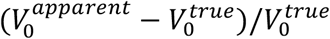.

Figure 6 considers the contribution of intravascular signal to the parameter estimates in Fig. 4 as a function of vessel radius. This contribution is generally small for R_2_′ and DBV at around ±1% for vessel radii greater than 10 μm. However, the intravascular signal appears to reflect a larger contribution when OEF is low, conditions where qBOLD contrast is low. Despite this the effect of the intravascular signal appears to be largely cancelled in the estimation of OEF (Fig. 6c,f). A reproduction of Fig. 4 without intravascular signal is included in the supplementary material for comparison and shows little discernible difference by eye (Fig. S6).

**Fig. 6.**
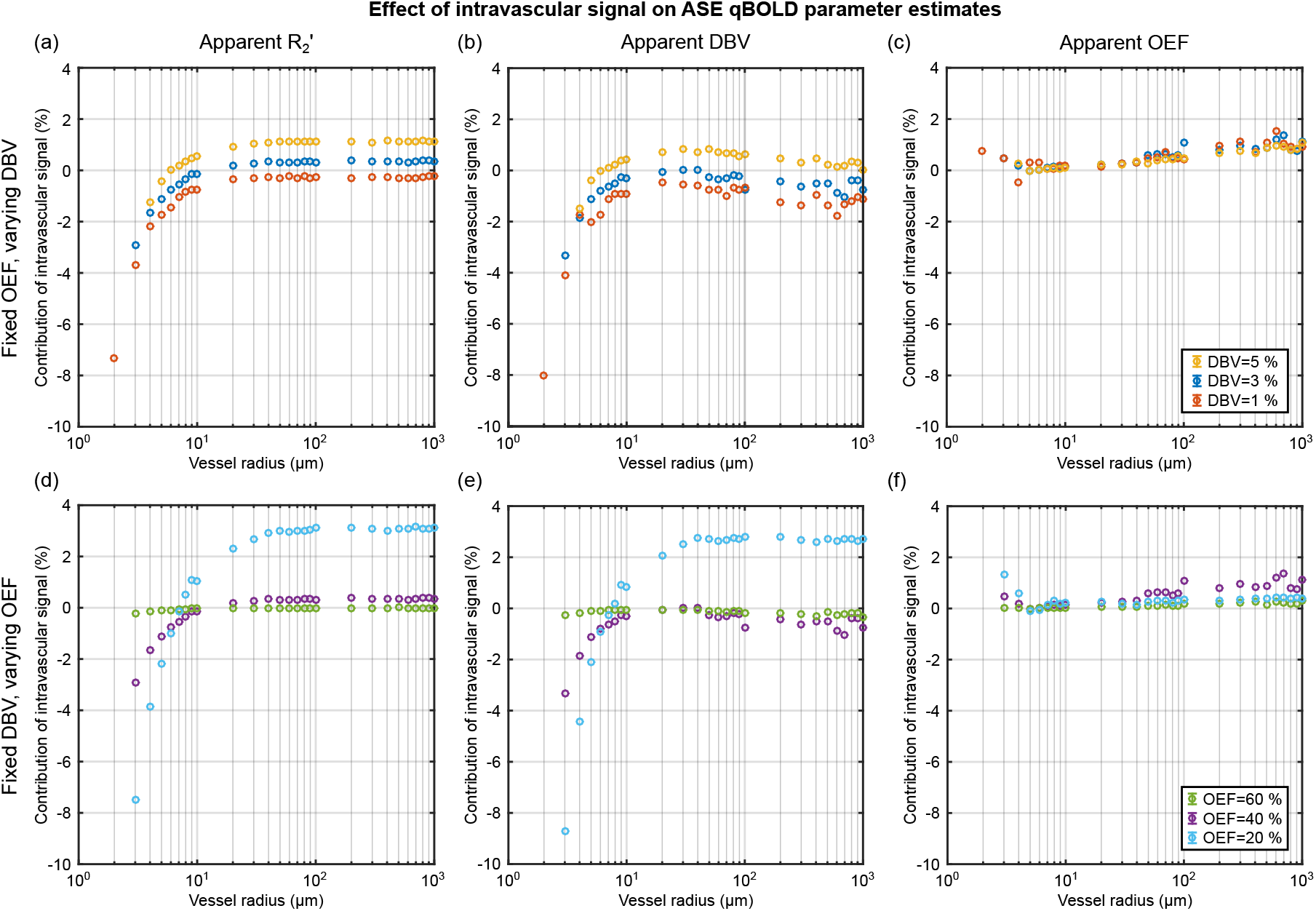
Investigation of the contribution of intravascular signal to qBOLD parameter estimates (PE) presented in Fig. 3. This contribution was quantified as the percentage difference between PEs simulated with and without intravascular signal i.e. (*PE*_*EV*_ − *PE*_*EV+IV*_)/*PE*_*EV+IV*_. The contribution of intravascular signal is observed to be relatively small for all parameters. Extravascular only PE results can be found in supplementary materials (Fig. S6).

Figure 7 investigates the origin of the DBV estimation error attributed either to an error in the measured signal at τ=0 (orange markers) or an error in the intercept extrapolated from long τ data (green markers). In the case of the former, 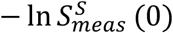 is plotted such that the sum of the two curves representing the apparent DBV (represented by grey shading). When interpreting these curves, it is useful to consider the orange markers as a reflection of the deviation of the spin echo from perfect refocusing (with positive values representing increased signal attenuation) and the green markers as a reflection of the deviation of the measured R_2_′ from the SDR qBOLD estimate of R_2_′. The former is found to be subject to increasing signal attenuation as vessel size is reduced, which is strongly affected by blood oxygenation via OEF. The latter is found to plateau and is relatively consistent with the SDR qBOLD model for vessel radii greater than approximately 20 μm.

**Fig. 7.**
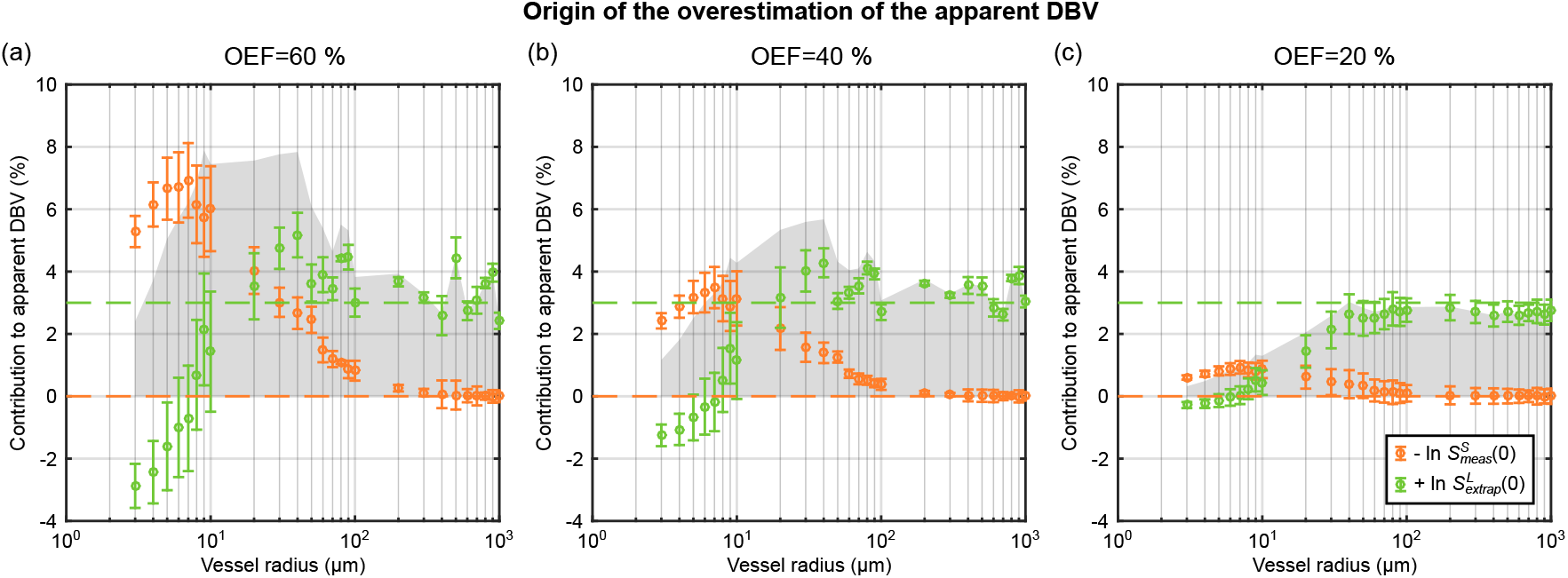
Investigation of the origin of the overestimation of the measured DBV for three different OEF values; (a) E_0_=60%, (b) E_0_=40%, (c) E_0_=20% (true V_0_=3%). The orange markers represent the natural log of the measured signal at τ=0, plotted here as 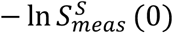, displaying increasing signal attenuation with decreasing vessel radius. Whilst the green markers represent the log of the intercept extrapolated from long τ data points 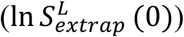 and appears more stable in the face of a reduced vessel radius. The sum of these curves is the apparent DBV as in Fig. 4 and represented here by the grey shaded area. Dashed lines display the prediction made by the SDR qBOLD model.

Figure 8 explores the combined effect of a distribution of vessel radii on parameter estimates from the SDR qBOLD model. The apparent R_2_′ is plotted against values of R_2_′ predicted by the SDR model via Eq. (2), with DBV estimated according to the working assumption described above (Fig. 8a). Data points are colour coded to reflect the true voxel deoxyhaemoglobin content in ml^dHb^/100 g^tissue^. A linear dependence is maintained, albeit with a shallower gradient than predicted by the SDR qBOLD model. A large amount of uncertainty is observed in estimates of apparent DBV over the large physiological range tested (Fig. 8b), with data points colour coded by true OEF value. However, this level of uncertainty does not propagate into estimates of apparent OEF (Fig 8c) where data points are colour coded by true DBV. Apparent OEF increases monotonically between 0 and 50%, but reaches a plateau for higher values, and is inappropriately scaled compared with the true OEF i.e. the full range of OEF is represented by apparent OEF values between 16% and 25%. In a similar manner to Fig. 5, the percentage error in the apparent DBV can be plotted as a function of true OEF (Fig. 9). As noted for the single vessel radius simulations, this error is strongly OEF dependent.

**Fig. 8.**
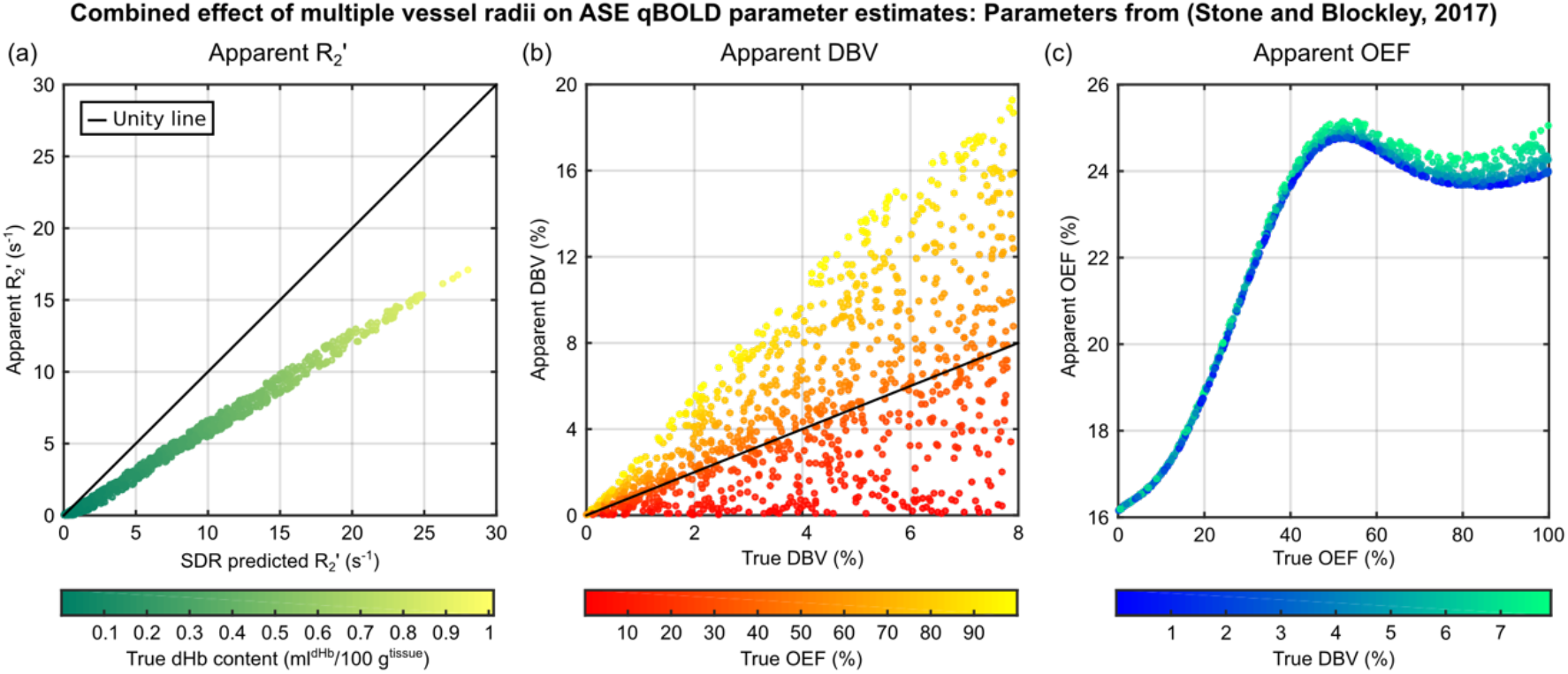
The effect of multiple vessel radii simulations on the qBOLD parameter estimates was considered by generating many pairs of OEF and CBV values. ASE pulse sequence parameters were t_E_=80 ms with τ=0 and τ=16 to 64 ms in 4 ms steps following the work of (Stone and Blockley, 2017). (a) The apparent R_2_′ is linearly dependent on the R_2_′ predicted by the SDR model, but with a different gradient. (b) A large amount of uncertainty in the apparent DBV is observed. (c) The apparent OEF appears to plateau beyond 50%, but monotonically increases with true OEF for lower values. Markers are coloured to reflect true dHb content, true OEF and true DBV for parts (a), (b) and (c), respectively.

**Fig. 9.**
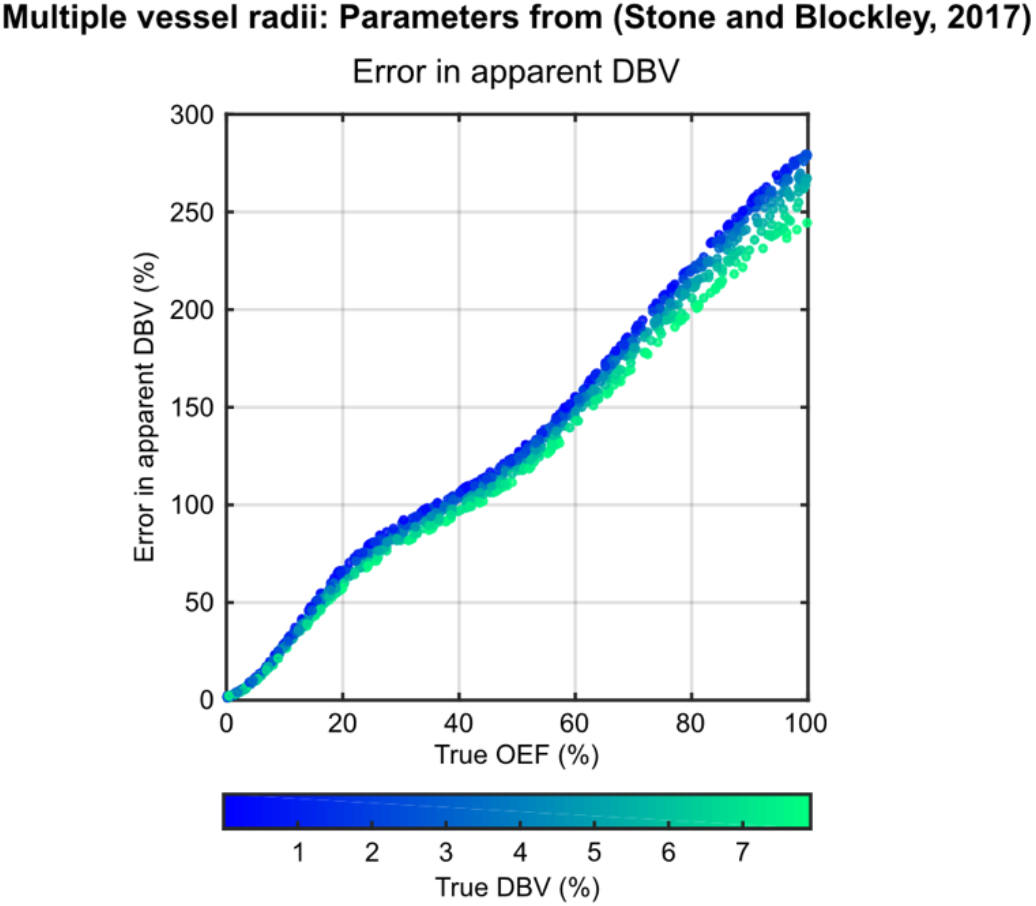
The uncertainty in DBV in Fig. 8 was investigated by plotting apparent DBV as a function of true OEF. ASE pulse sequence parameters are the same as detailed in Fig. 8. The results suggest that the error in the apparent DBV is OEF dependent. Markers are coloured to reflect their true DBV.

These simulations were repeated for different ASE pulse sequence parameters, namely variations in t_E_ and τ, and included in supplemental material. The results in Fig. S7 largely mirror those in Fig. 8 with the following variations. The slope of the relationship between apparent R_2_′ and SDR qBOLD predicted R_2_′ is slightly reduced for the alternative parameters (Fig. S7a). More noticeable is the reduction in the range of apparent DBV values (Fig. S7b), with the error in the apparent DBV reduced by more than a half (Fig. S8). Whilst the apparent OEF is also inappropriately scaled, the relationship with true OEF is more monotonic in nature.

## Discussion

In this study numerical simulations were used to investigate the effect of diffusion on ASE based qBOLD measurements and the origin of DBV overestimation in such measurements. In contrast to the previously observed shift of the GESSE signal maximum due to the effect of diffusion, the ASE signal was observed to maintain its symmetry as vessel radius is reduced and the effect of diffusion is increased. Two hypotheses for the origin of the observed DBV overestimation were tested: (i) the effect of intravascular blood signal and (ii) the effect of diffusion on the extravascular tissue signal. The presence of intravascular blood signal was found to have a minor effect on qBOLD parameter estimates. It is therefore unlikely to be responsible for the majority of the overestimation observed in DBV measurements. In contrast, the extravascular signal was shown to have a very strong dependence on vessel radius providing the potential for a large error in DBV and is considered to be the dominant cause of DBV overestimation. Furthermore, the error in DBV is predicted to be blood oxygen saturation level dependent. Integration of these single vessel radius simulations via a more physiologically realistic vessel distribution revealed three main findings. Firstly, that the relationship between the apparent R_2_′ and deoxyhaemoglobin content is retained. Secondly, there is an inherent uncertainty in estimates of DBV. Finally, this uncertainty is not propagated to apparent OEF estimates, but results in inappropriate scaling of these estimates. Furthermore, the monotonic behaviour of the relationship between apparent and true OEF was found to be dependent on the pulse sequence parameters t_E_ and τ. These results provide new directions for improving the modelling of ASE qBOLD signal and the reduction of systematic error in parameter estimates of OEF and DBV.

### Effect of diffusion on ASE measurements

Whilst several studies have investigated the qBOLD signal as acquired by the GESSE pulse sequence (Christen et al., 2014; Dickson et al., 2011, 2010; Pannetier et al., 2014), this study considered whether the signal decay under an ASE acquisition behaves in the same way. One particular characteristic of the GESSE pulse sequence concerns the maximum of the qBOLD signal decay curve. This would ordinarily be expected to coincide with the spin echo (τ= 0), but has been shown to be shifted towards negative τ values (Fig. S5a) for the GESSE sequence in the presence of diffusion (Dickson et al., 2010; Pannetier et al., 2014). However, this effect is not observed in simulations of the extravascular signal acquired using an ASE pulse sequence (Fig. 3a), where the signal maximum was found to be close to the spin echo (τ=0). However, the GESSE and ASE sequences differ in an important way. The t_E_ of each successive τ value increases in the GESSE experiment and hence the time for protons to diffuse around blood vessels increases. Whilst the tE is constant for all τ values in the ASE method and hence the time for diffusion is also constant. This would suggest that there is a t_E_ dependent component of the R_2_′-weighted signal decay. Such a component has previously been included as a correction to estimates of R_2_′ (Berman et al., 2017).

This study also considered the R_2_′-weighted contribution of the blood to the qBOLD signal using a recently proposed model (Berman and Pike, 2018). In common with the extravascular results, the ASE blood signal is symmetric with respect to the spin echo, but decays far less as a function of τ (Fig. 3b). However, the signal is heavily attenuated at all τ values compared with the extravascular simulations. This is in contrast to simulations of the GESSE blood signal, which are highly shifted to negative τ values and present largely as an exponential decay (Fig. S5b).

### Origin of DBV overestimation

Simulations of the combined intravascular and extravascular signal revealed a vessel radius dependent overestimation of DBV for vessel radii greater than 5 μm (Fig. 4b,e). The error in the apparent DBV was found to be OEF dependent (Fig. 5). However, at larger radii (approaching 1 mm) estimates of DBV were consistent with ground truth values. The contribution of intravascular signal to these parameter estimates was determined by comparing simulations with (Fig. 4) and without (Fig. S6) an intravascular compartment. A small and largely vessel radius independent effect (for R_c_>10 μm) was observed (Fig. 6b,e). The effect of intravascular signal was more pronounced for smaller vessel radii and low OEF, where the relative contribution of intravascular signal is increased by weak extravascular contrast. Despite this, the overestimation of DBV is dominated by the effect of diffusion on the extravascular signal.

Finally, Eq. (4) provides the opportunity to consider whether the systematic error in DBV originates in the measurement of the spin echo 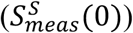, the intercept extrapolated from the long τ regime 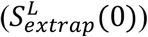 or a combination of both, as explored in Fig. 7. For vessel radii greater than 20 μm additional signal attenuation of 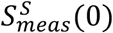 is the main driver of overestimation of DBV. However, for vessels with radii below 20 μm, errors in 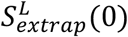 provide an additional confound to DBV estimation. These results are consistent with the characteristics of gradient echo versus spin echo BOLD vessel size sensitivity, which correspond to 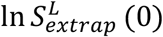 and 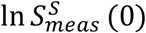, respectively (Boxerman et al., 1995). For the smallest vessel radii the apparent R_2_′ is reduced relative to the value expected by the SDR qBOLD model (Fig. 4a,d) due to diffusional narrowing, such that 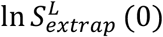 is also reduced (Fig. 7). Similarly, additional unrecoverable signal decay due to diffusion narrowing results in a decrease in the value of 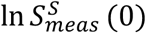, which is analogous to an increase in apparent R_2_ and is strongest for capillary sized vessels (Note that Fig. 7 plots 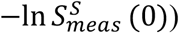. With increasing vessel radius, R_2_′ approaches the SDR qBOLD model prediction and the value of 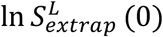 approaches a constant value. Similarly the attenuation of the spin echo is reduced as the SDR is approached and 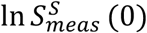 reaches its minimum. Therefore, when the differing profiles of these phenomena are combined the form of the apparent DBV as a function of vessel radius can be described.

### Effect of a physiological vessel radius distribution

Having established the vessel radius dependence of the qBOLD signal, the implications for experimental measurements were considered. In order to integrate the single vessel radius results, a vessel distribution with a small number of discrete vessel radii was selected. This enabled different oxygenation levels to be associated with different vessel types. A wide physiological range was investigated by randomly selecting pairs of OEF and CBV values. The apparent R_2_′ was found to be tightly correlated with the R_2_′ predicted by SDR qBOLD model (Fig. 8a). This is important as it demonstrates that the relationship between R_2_′ and the voxel deoxyhaemoglobin content (proportional to the product of deoxyhaemoglobin concentration and DBV) is maintained despite the effects of diffusion. It should therefore be possible to quantify maps of R_2_′ in terms of deoxyhaemoglobin content with appropriate scaling. Likewise with improved quantification of DBV, either through improvements to the qBOLD technique or via an additional experimental technique (Blockley et al., 2013; Lee et al., 2018), accurate measurements of OEF are possible. A large amount of uncertainty was observed in the apparent DBV (Fig. 8b). This was demonstrated to be blood oxygenation dependent i.e. a function of OEF (Fig. 9). This is consistent with the results of the single vessel simulations (Fig. 5) and demonstrates the important contribution of smaller vessel radii. This also explains why this uncertainty does not propagate into the apparent OEF, since the percentage error in apparent DBV is constant at each OEF level (Fig. 8c). However, the increasing percentage error in apparent DBV with OEF (Fig. 9) results in a progressive underestimation of apparent OEF. A plateau in the apparent OEF limits the maximum measured OEF to approximately 50%. Despite this the remaining range covers the majority of the expected healthy physiological range (Marchal et al., 1992). These simulation were repeated for an alternative set of ASE pulse sequence parameters, replicating the effects observed for R_2_′ and DBV (Fig S7a,b and Fig. S8). A monotonic relationship between apparent and true DBV was revealed and although the linear portion is limited to the range between 20% and 80% this encompasses the range reported in ischaemic stroke lesions defined using diffusion weighted imaging (Guadagno et al., 2006). The underlying mechanisms for this altered behaviour are inherently multidimensional and require further systematic investigation. However, these results demonstrate that there is additional scope for optimisation of qBOLD through changes to t_E_ and the range of τ values.

Finally, the results of these multi-radius simulations appear to be consistent with previous measurements of OEF=21±2% and DBV=3.6±0.4% (Stone and Blockley, 2017). Under the assumption that a true OEF of 40% is healthy, Fig. 8 would predict an apparent OEF of 24%. Likewise Fig. 9 would predict the percentage error in the apparent DBV is 100%, which would reduce the measured value above to 1.8%. This would bring these measurements in line with other MR based measurements of DBV at 1.75% (He and Yablonskiy, 2007) and venous CBV at 2.2% (Blockley et al., 2013). For the alternative ASE pulse sequence parameters Fig. S7 predicts an apparent OEF of 40% for a true OEF value of 40%, which is consistent with experiments (An and Lin, 2003). However, Fig. S7 also predicts that the dynamic range of OEF is compressed, suggesting that modulations of OEF with respect to this baseline would be underestimated.

### Limitations

Whilst the simulation methodology used in this study has identified some limitations of the current implementation of ASE based qBOLD, it also offers an opportunity to optimise future implementations. Further simulations could be used to identify optimal values of t_E_ and τ which maximise the linearity of the relationship between apparent OEF and the ground truth. They could also be used to estimate a more appropriate scale factor for OEF estimation by treating the 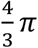 geometry factor in Eq. (2) as an arbitrary scale factor. Such an approach has previously been used in calibrated BOLD to great effect (Griffeth and Buxton, 2011).

The results of this study rely on a detailed model of the qBOLD signal. However, in this implementation it only accounts for the intra- and extravascular signal contributions of a single distribution of blood vessels in grey matter. Whilst this two compartment model was sufficient to investigate the origin of DBV overestimation in ASE based qBOLD, a more realistic model might include the signal contributions of cerebral spinal fluid (Dickson et al., 2009), the myelin in white matter (Bouvier et al., 2013), desaturated arterial blood vessels (Boas et al., 2008), the effect of iron deposition (Wismer et al., 1988) or different vessel radius distributions (Germuska et al., 2013; Lauwers et al., 2008). As such these contributions to the qBOLD signal may also provide fertile ground for future exploration.

In addition, this study did not consider the effects of magnetic field inhomogeneity or noise on the measured signal. The former has been extensively studied experimentally (Blockley and Stone, 2016; Dickson et al., 2010; Yablonskiy, 1998), but may benefit from more detailed simulations to test the assumptions of these correction schemes. The latter poses a particular problem for the analysis approach described by Eq. (16) and (17), is reliant on a single measurement in the short τ regime acquired at the spin echo. A broader range of measurements in the short τ regime could be incorporated into the analysis using a non-linear model fitting approach based on Eq. (3) and (4), which may also result in reduced uncertainty in parameter estimates. Further improvements could be achieved by using a more sophisticated analysis approach, such as a Bayesian framework which would enable prior knowledge about physiological parameters to be incorporated (Chappell et al., 2008). Finally, this study has demonstrated that by altering the ASE acquisition parameters it is possible to address some of the limitations of our existing experimental approach. Optimisation of these parameters will form the focus of future work.

## Conclusion

The ASE qBOLD signal decay was found to be symmetric with respect to the spin echo, in contrast to previous simulation of the GESSE pulse sequence. Overestimation of DBV by ASE based qBOLD was found to be dominated by the effect of diffusion on extravascular signal decay, with the presence of intravascular blood signal having only a small contribution. Integrating the results of single vessel simulations using an *in vivo* distribution of vessel radii revealed several limitations of current measurements and provides a foundation for future optimisation of ASE based qBOLD acquisitions.

## Funding Acknowledgements

This work was supported by the Engineering and Physical Sciences Research Council [grant number EP/K025716/1]. The Wellcome Centre for Integrative Neuroimaging is supported by core funding from the Wellcome Trust [grant 203139/Z/16/Z].

## Appendix A

The results of the simulations performed in this study can be accessed via the Oxford University Research Archive, doi: https://doi.org/10.5287/bodleian:mvPY99a9D. Furthermore, the code used to generate these simulation results and to analyse experimental data can be downloaded from the Zenodo repository, doi: https://doi.org/10.5281/zenodo.3241420. Supplementary data related to this article can be found at <DOI>.

---

**Fig. S1.**
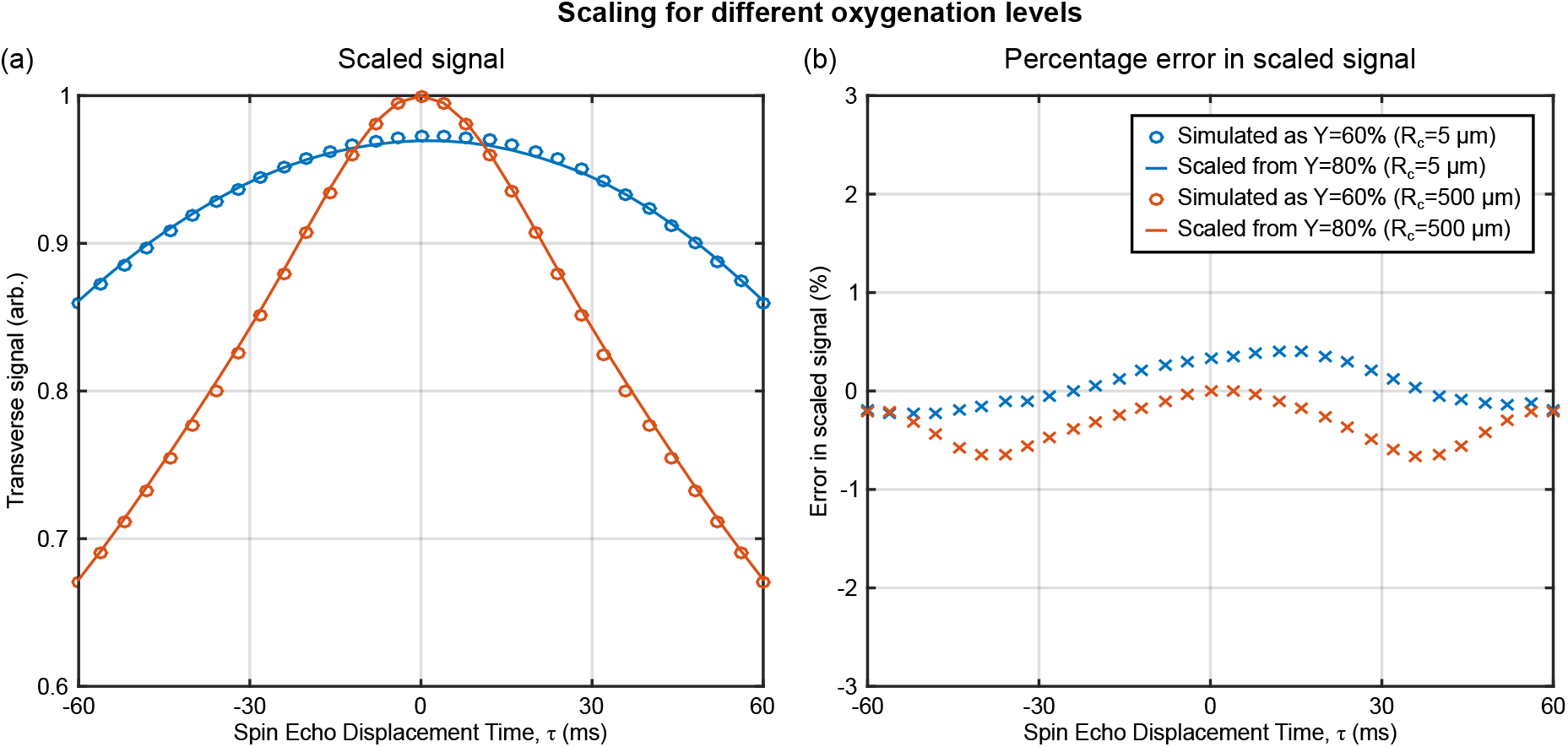
Monte Carlo simulations can be accelerated to calculate different oxygenations by saving the phase accumulated by each proton for a nominal oxygenation value. Since phase scales linearly with oxygenation, the simulated phase accrual can be scaled to a target oxygenation prior to the estimation of the signal (Blockley et al., 2008). (a) Two examples are shown here: (i) a vessel radius of 5 μm simulated as Y=60% (blue markers) and simulated as Y=80% and scaled to Y=60% (blue line) and (ii) a vessel radius of 500 μm simulated as Y=60% (red markers) and simulated as Y=80% and scaled to Y=60% (red line). (b) The percentage error between the simulated and scaled signals are also displayed i.e. (*S*^*simulated*^ − *S*^*scaled*^)/*S*^*simulated*^.

**Fig. S2.**
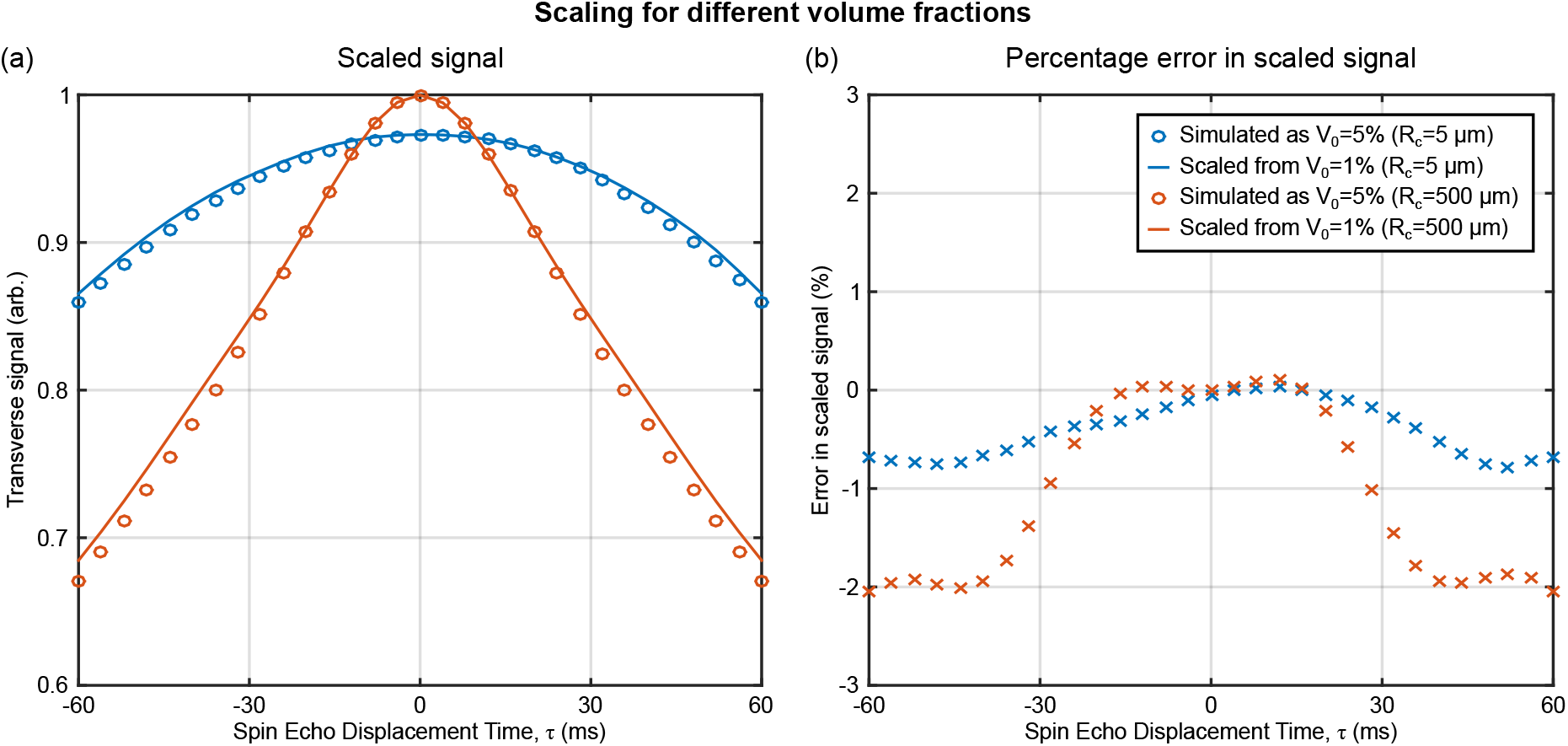
Monte Carlo simulations can also be accelerated to calculate different blood volume fractions. This is achieved by estimating an oxygenation and vessel radius dependent shape function (Dickson et al., 2011; Kiselev and Posse, 1999). This shape function can then be arbitrarily scaled for different blood volume fractions. (a) Two examples are shown here: (i) a vessel radius of 5 μm simulated as V_0_=5% (blue markers) and simulated as V_0_=1% and scaled to V_0_=5% (blue line) and (ii) a vessel radius of 500 μm simulated as V_0_=5% (red markers) and simulated as V_0_=1% and scaled to V_0_=5% (red line). (b) The percentage error between the simulated and scaled signals are also displayed i.e. (*S*^*simulated*^ − *S*^*scaled*^)/*S*^*simulated*^.

**Fig. S3.**
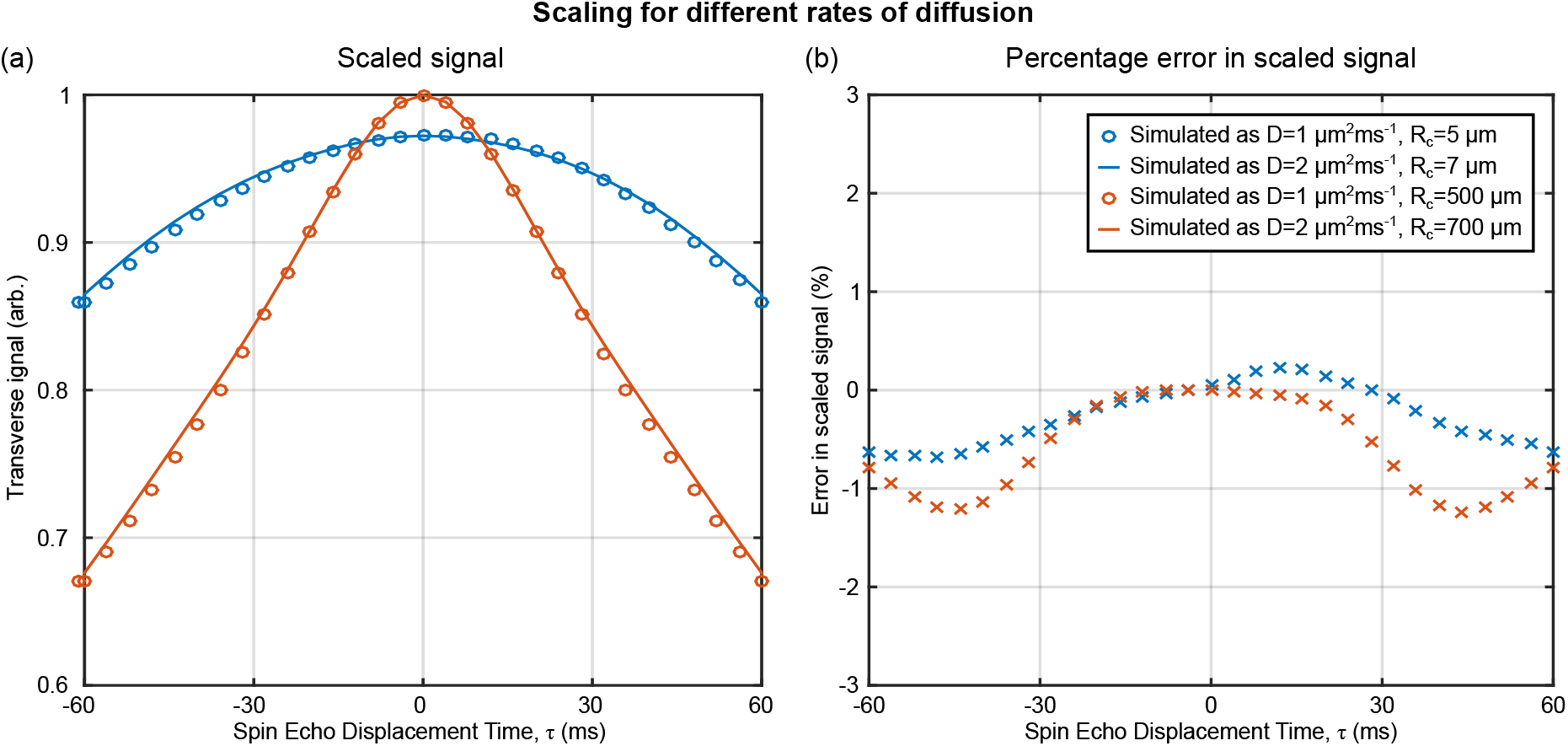
Monte Carlo simulations can also be accelerated to simulate the effect of different diffusion coefficients, D. The effect of diffusion is a function of vessel radius and diffusion coefficient through the characteristic diffusion time 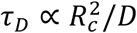 (Yablonskiy and Haacke, 1994). Therefore, the effect of a change in diffusion can be simulated by scaling the vessel radius e.g. scaling D by a factor of 2 requires R_c_ to be scaled by 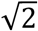. (a) Two examples are shown here: (i) a vessel radius of 5 μm simulated with D=1 μm^2^ms^−1^ (blue markers) compared with a vessel radius of 7 μm simulated with D=2 μm^2^ms^−1^ (blue line) and (ii) a vessel radius of 500 μm simulated with D=1 μm^2^ms^−1^ (red markers) compared with a vessel radius of 700 μm simulated with D=2 μm^2^ms^−1^ (red line). (b) The percentage error between the simulated and scaled signals are also displayed i.e. (*S*^*simulated*^ − *S*^*scaled*^)/*S*^*simulated*^.

**Fig. S4.**
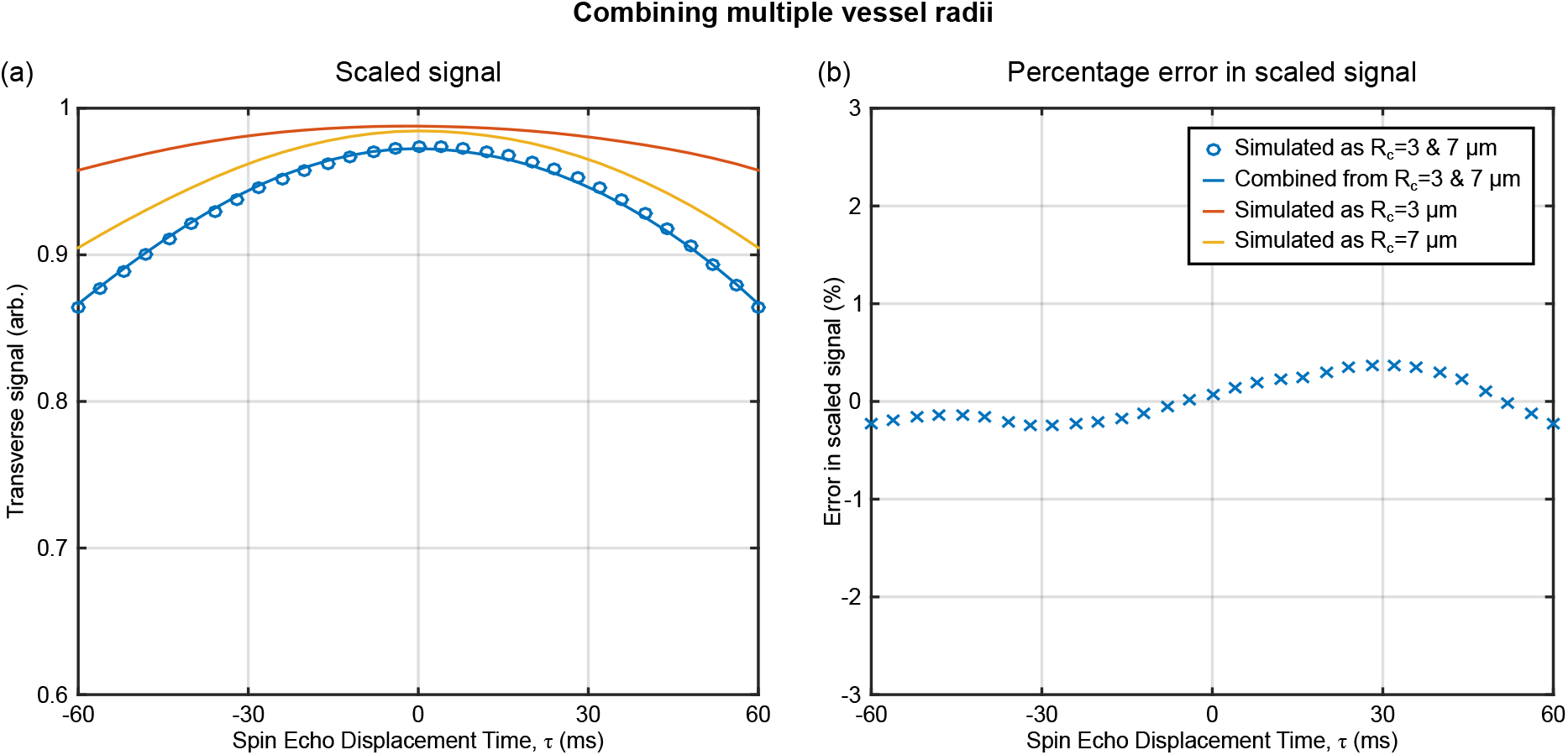
The simulation of systems with multiple vessel radii can be simulated by combining the results of multiple single vessel radius simulations (Dickson et al., 2011; Kiselev and Posse, 1999). (a) In this example, simulations of a system with V_0_=5% equally split between vessels with R_c_=3 μm and R_c_=7 μm (blue markers). Single vessel simulations of R_c_=3 μm (red line) and R_c_=7 μm (yellow line) with V_0_=2.5%. The combined effect of these single vessel simulations is obtained by taking the product of these signal curves (blue line). (b) The percentage error between the simulated and combined signals are also displayed i.e. (*S*^*simulated*^ − *S*^*scaled*^)/*S*^*simulated*^.

**Fig. S5.**
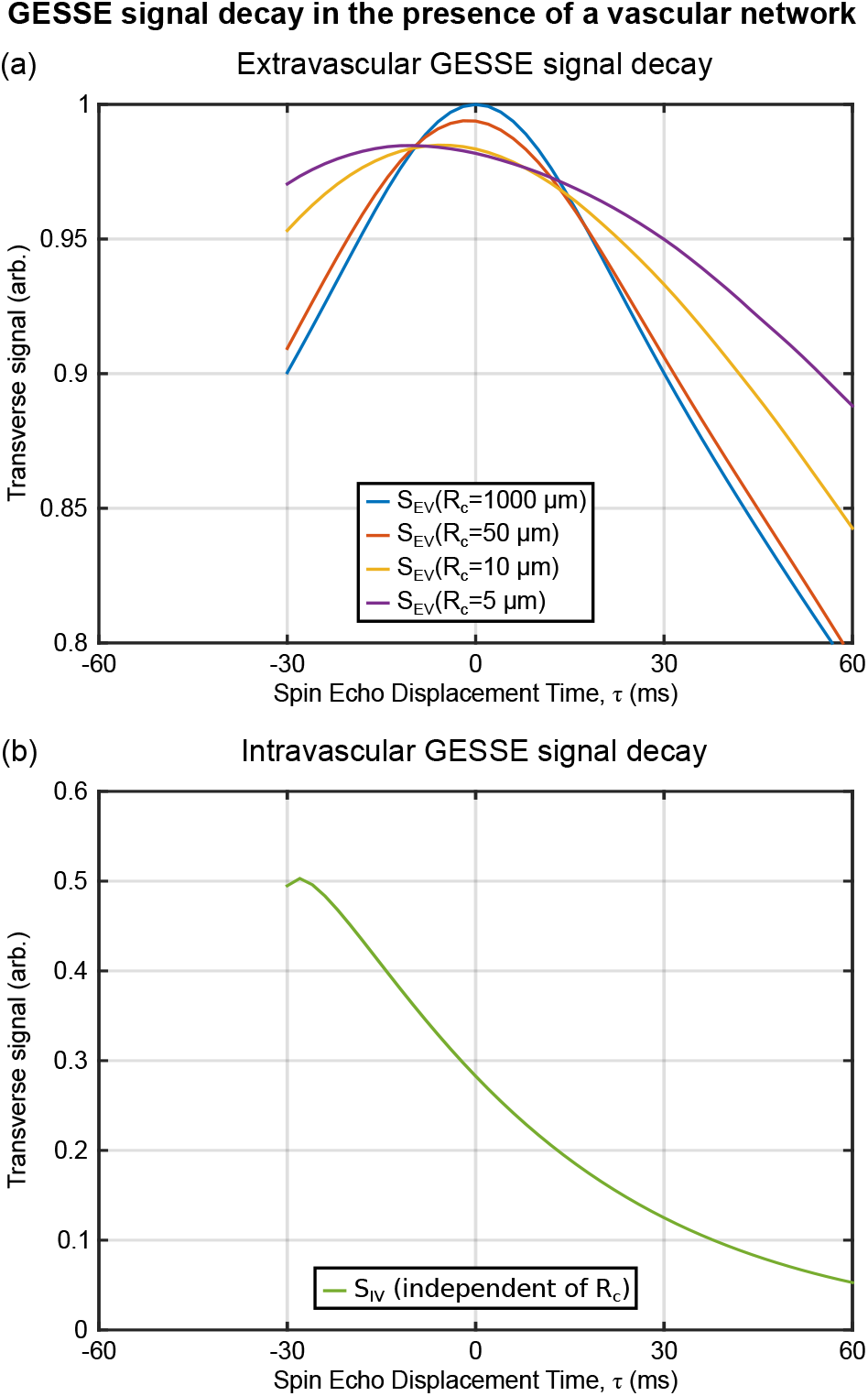
Examples of the signal decay from the GESSE pulse sequence. The extravascular signal (S_EV_) decay is observed to be asymmetric with respect to τ=0 as the vessel radius is reduced. (b) The intravascular signal (S_IV_) decay shows considerable signal attenuation which is highly asymmetric with respect to τ and appears almost exponential in form.

**Fig. S6.**
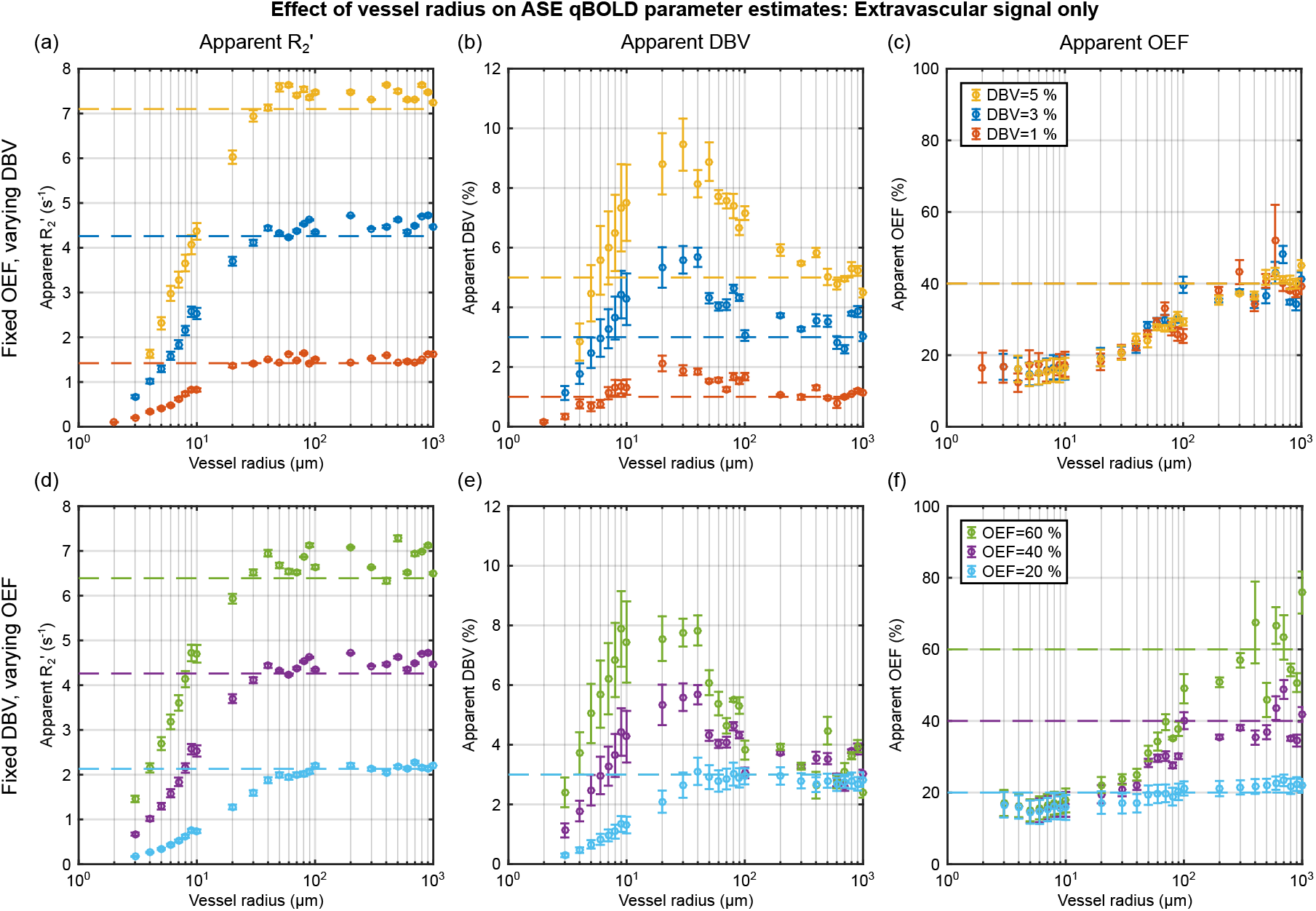
Reproduction of Fig. 3 for simulations of the extravascular signal only. As in Fig. 3, simulations were first performed with a fixed OEF (E_0_=40%) three DBV values (top) then with a fixed DBV (V_0_=3%) and three values of OEF (bottom). The apparent R_2_′ (left) is estimated for each OEF-DBV pair and presented alongside the R_2_′ values predicted by the SDR qBOLD model (dashed lines). Likewise the apparent DBV (centre) and apparent OEF (right) are presented alongside the true DBV and OEF, respectively, (dashed lines).

**Fig. S7.**
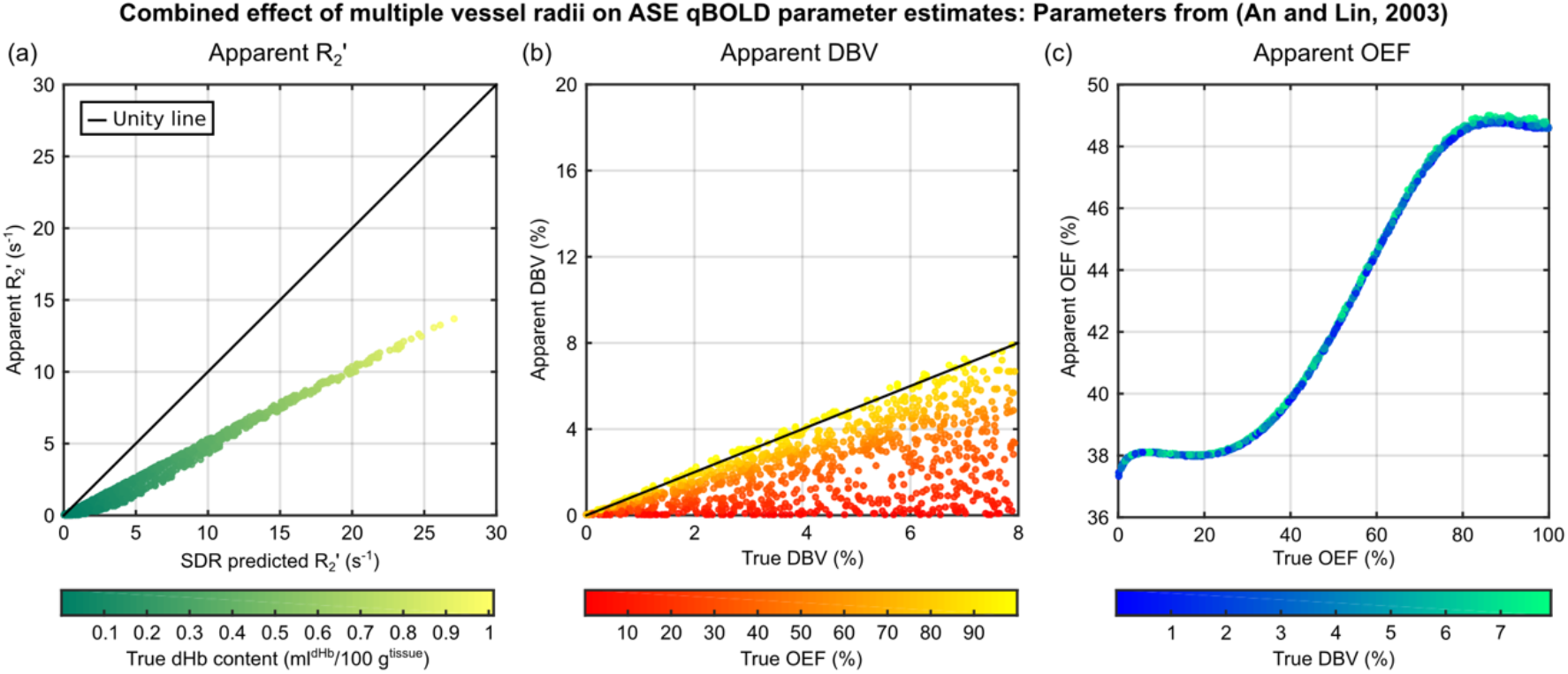
The effect of multiple vessel radii simulations on the qBOLD parameter estimates was considered by generating many pairs of OEF and CBV values. ASE pulse sequence parameters were tE=64 ms with τ=0 and τ=10 to 18 ms in 4 ms steps following the work of (An and Lin, 2003). (a) The apparent R_2_′ is linearly dependent on the R_2_′ predicted by the SDR model, in common with the parameters used in Fig. 7 but with a shallower gradient. (b) Uncertainty is similarly observed in the apparent DBV, but with a reduced range of values. (c) In contrast to Fig. 7c, the apparent OEF has a largely monotonic relationship with the true OEF. Markers are coloured to reflect true dHb content, true OEF and true DBV for parts (a), (b) and (c), respectively.

**Fig. S8.**
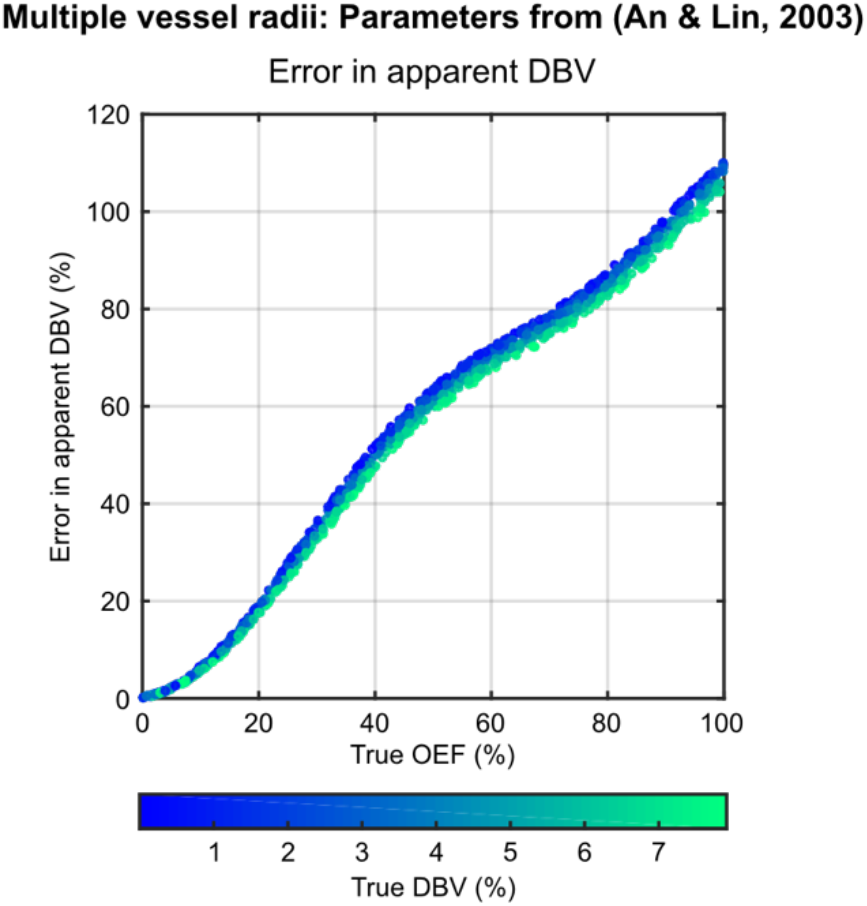
The uncertainty in DBV in Fig. S7 was investigated by plotting apparent DBV as a function of true OEF. ASE pulse sequence parameters are the same as detailed in Fig. S7. In common with Fig. 9, these results suggest that the error in the apparent DBV is OEF dependent. Markers are coloured to reflect their true DBV.

